# Cortical dissociation of spatial reference frames during place navigation

**DOI:** 10.1101/2025.06.25.661569

**Authors:** Patrick A. LaChance, Elaina R. Gross, Shruthi Sankaranarayanan, Michael E. Hasselmo

## Abstract

Animals rely on both sensory perception and memory when navigating relative to learned allocentric locations. Incoming sensory stimuli, which arrive from an egocentric perspective, must be integrated into an allocentric reference frame to allow neural computations that direct an animal toward a learned goal. This egocentric-allocentric spatial transformation has been proposed to involve projections from the rodent postrhinal cortex (POR), which receives strong visual input, to the medial entorhinal cortex (MEC), which contains allocentric spatial cell types such as grid and border cells. A major step toward understanding this transformation is to identify how POR and MEC spatial representations differ during place navigation, which is currently unknown. To answer this question, we recorded single neurons from POR and MEC as rats engaged in a navigation task that required them to repeatedly visit a learned uncued allocentric location in an open field arena to receive a randomly scattered food reward. While neurons in both regions displayed strong tuning to the spatial structure of the environment, neither showed bias toward the goal location despite strongly biased behavior. Critically, when local visual landmarks were manipulated to place the visual scene in conflict with the learned location, POR neurons adjusted their tuning preferences to follow the visual landmarks, while MEC neurons remained in register with the true global reference frame. These findings reveal a strong dissociation between POR and MEC spatial reference frames during place navigation and raise questions regarding the mechanisms underlying integration of POR egocentric signals into the MEC allocentric spatial map.

## Introduction

Success in the wild often depends on remembering and navigating relative to specific locations that are not immediately perceptible. For example, an animal may need to execute a path from its home base to an unmarked food cache. This process requires the animal to interface incoming sensory stimuli, such as salient visual landmarks, with a memory of how the goal location is situated relative to those stimuli. The initial perception of spatial landmarks is considered ‘egocentric’ because it occurs from the animal’s first-person perspective, while the goal location itself is considered ‘allocentric’ because it is defined with respect to the objective spatial layout of the world (Gallistel, 1990; Klatzky, 1998). Exactly how these unique processing streams combine in the brain remains elusive.

Allocentric spatial coding in the brain during navigation has been extensively investigated with respect to the hippocampus, which appears critical for supporting allocentric spatial reference memory (Morris et al., 1982; Packard and McGaugh, 1996) and contains place cells that encode specific allocentric locations (O’Keefe, 1976). The hippocampus is heavily innervated by the entorhinal cortex (Steward and Scoville, 1976; Witter et al., 2000), whose medial subdivision (MEC) contains a variety of cell types that code for space in an allocentric reference frame, such as grid, border, and head direction (HD) cells (Hafting et al., 2005; Sargolini et al., 2006; Savelli et al., 2008; Solstad et al., 2008; Boccara et al., 2010). Importantly, sensory input is thought to reach MEC extensively via projections from the postrhinal cortex (POR) (Naber et al., 1997; Burwell and Amaral, 1998a; Witter et al., 2000; Koganezawa et al., 2015), which receives heavy visual input (Burwell and Amaral, 1998b; Burwell, 2000; Tomás Pereira et al., 2016; Beltramo and Scanziani, 2019) and contains neurons primarily tuned to the egocentric layout of the landmarks and geometry of the surrounding environment (Gofman et al., 2019; LaChance et al., 2019; LaChance et al., 2022; LaChance and Taube, 2023b). While this dissociation between POR egocentric and MEC allocentric reference frames has been demonstrated during random foraging experiments (LaChance et al., 2019; LaChance et al., 2022; LaChance and Hasselmo, 2024), it remains unknown how POR and MEC may differentially encode aspects of the environment during navigation relative to a learned allocentric place.

## Results

### Rats quickly learn a place navigation task

We trained rats (n = 4) to repeatedly visit an uncued 15 x 15 cm goal zone within a 120 x 120 cm open field with a single orienting visual landmark (large white ‘cue card’) placed along the south wall (Fig. 1A). Entry into the goal zone, detected by an overhead camera, triggered an overhead dispenser to release sucrose pellets that randomly scattered throughout the open field, encouraging the animals to sample the entire task environment. The goal zone always occupied the same location in the northwest quadrant of the environment for all animals, centered 35 cm from the west wall and 45 cm from the north wall (Fig. 1A). After triggering a reward, a 10 second reward timeout period prevented the rats from immediately receiving another reward delivery. Note that this task is similar to the place preference paradigm used to test goal coding by place cells (Rossier et al., 2000; Hok et al., 2007b), and differs only in that the animals were not required to wait in the goal zone to receive reward.

**Figure 1.**
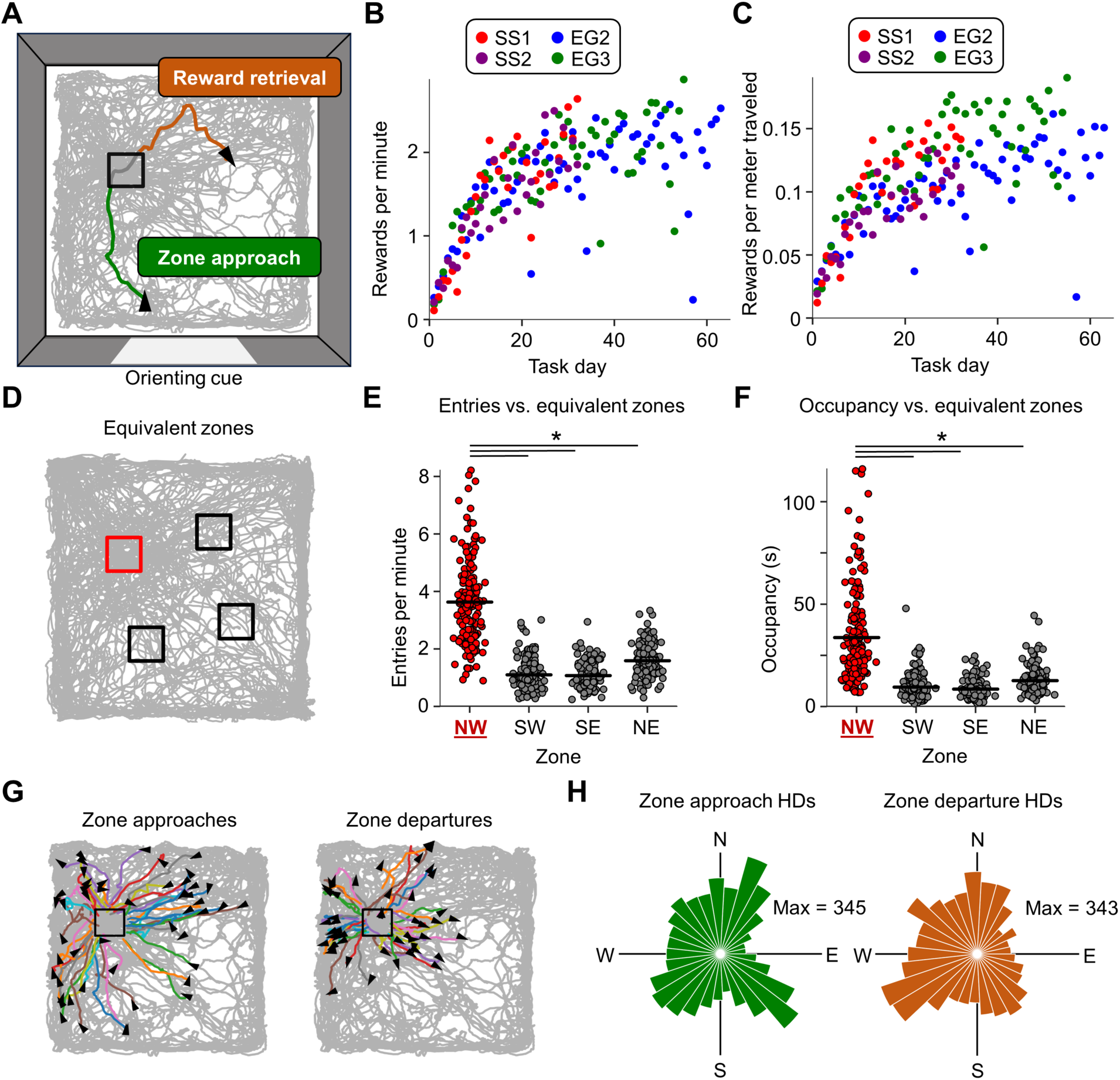
Place navigation task and performance. **A)** Top-down illustration of the place navigation task. The gray trace indicates the animal’s path over the entire task session, the black box indicates the location of the goal zone, the green line indicates the animal’s path during one approach to the goal (“zone approach”; 1 second trajectory), and the orange line indicates the animal’s path after exiting the zone (“reward retrieval”; 3 second trajectory). Black triangles indicate the animal’s head direction at the start of the zone approach and end of the reward retrieval trajectories. An orienting cue (white card) is shown along the south wall of the enclosure. **B)** Learning curve for all animals during standard task sessions performed each day, indicating the number of rewards triggered per minute across all task days. **C)** Same as **(B)** but indicating the number of rewards triggered per meter traveled. **D)** Example trajectory plot showing the true goal zone (red) as well as rotationally equivalent zones (black) that were used for comparison. **E)** Strip plot indicating, for all standard task sessions across all animals, the number of entries per minute into the goal zone vs. rotationally equivalent zones (repeated measures ANOVA, *F*(3, 483) = 358.85, *P* = 6.23e-66; paired *t*-tests, NW vs. SW, *t*(161) = 22.15, *P* = 2.93e-50; NW vs. SE, *t*(161) = 20.99, *P* = 1.84e-47; NW vs. NE, *t*(161) = 18.05, *P* = 4.70e-40). **F)** Same as **(E)** but for time spent in the goal zone vs. rotationally equivalent zones. (repeated measures ANOVA, *F*(3, 483) = 158.32, *P* = 3.77e-33; paired *t*-tests, NW vs. SW, *t*(161) = 13.31, *P* = 2.86e-27; NW vs. SE, *t*(161) = 13.91, *P* = 6.67e-29; NW vs. NE, *t*(161) = 12.18, *P* = 3.92e-24). Note that animals had more entries per minute and higher occupancy in the goal zone compared to equivalent zones. **G)** Trajectory plots for an example standard task session (same as session shown in **(A)**) with overlaid trajectories indicating the animal’s path prior to zone entry (*left*) and following zone exit (*right*; 1 second trajectories). Note that approaches and departures covered the full range of directions. **H)** Polar histograms indicating the distribution of animal head directions during zone approaches (*left*) and zone departures (*right*) for all standard task sessions across all animals.

Rats quickly learned this task, reaching an average of 1 reward/minute within 10 training days (Fig. 1B, C), at which point neural recordings began. To assess behavioral bias toward the goal zone after the initial training period, we calculated the number of goal zone entries and time spent within the goal zone over the course of each session and compared them to rotationally equivalent zones in the other three quadrants of the environment (Fig. 1D). Rats showed significant behavioral bias toward the goal zone compared to equivalent zones in terms of both number of entries and occupancy time (Fig. 1E, F; Fig. S1), indicating that they had learned the location of the goal zone. Importantly, rats approached the goal zone from a relatively uniform distribution of angles (approach Rayleigh *r*: 0.046, departure Rayleigh *r*: 0.13; Fig. 1G, H; Fig. S1), suggesting that they had learned to approach a place (allocentric strategy) instead of recreating the sensory stimuli that accompany a particular path (egocentric strategy) which would change given different approach directions.

### POR and MEC/PaS neurons uniformly encode the task space during place navigation

Recording of neural data began once animals reached a rate of approximately 1 reward/minute (Fig. 1B). In total, 415 neurons were recorded from POR (all four animals) and 286 were recorded from MEC or the adjacent parasubiculum (MEC/PaS; three of the animals; Fig. 2A; Table 1) during the standard place navigation task. For POR cells, we focused on cells with two major cell types: center-bearing (CB) cells, which fire preferentially when the boundaries or center of the environment fall at a specific angle relative to the animal’s heading (i.e., front/back/left/right; Gofman et al., 2019; LaChance et al., 2019); and head direction (HD) cells, which fire when the animal faces a certain direction in allocentric coordinates (i.e., north/south/east/west; Fig. 2B, C; LaChance et al., 2022). In MEC/PaS, we focused on grid cells, which fire at an array of allocentric locations that fall at the vertices of a triangular lattice (Hafting et al., 2005); border cells, which fire preferentially along one wall of the environment (Barry et al., 2006; Solstad et al., 2008; Lever et al., 2009); non-grid spatial cells, which possess allocentric location preferences that do not form a grid or border pattern (Diehl et al., 2017); as well as HD cells (Sargolini et al., 2006). Note that POR HD cells have been previously demonstrated to combine allocentric directional information with egocentric perception of visual landmarks (LaChance et al., 2022), becoming bidirectionally tuned when visual cues are made bidirectionally symmetrical, which distinguishes them from more ‘classic’, purely unidirectional HD cells commonly found elsewhere in the brain (Taube et al., 1990; Clark et al., 2024). For this reason, we will refer to POR HD-tuned neurons as landmark modulated-HD (LM-HD) cells. In general, MEC/PaS neurons tended to have higher HD mean vector lengths (MVLs; similar to ‘classic’ HD cells), lower CB MVLs, and higher spatial information content than POR neurons, along with including grid cells and theta-rhythmic cells which were largely absent in POR (Fig. S2).

**Figure 2.**
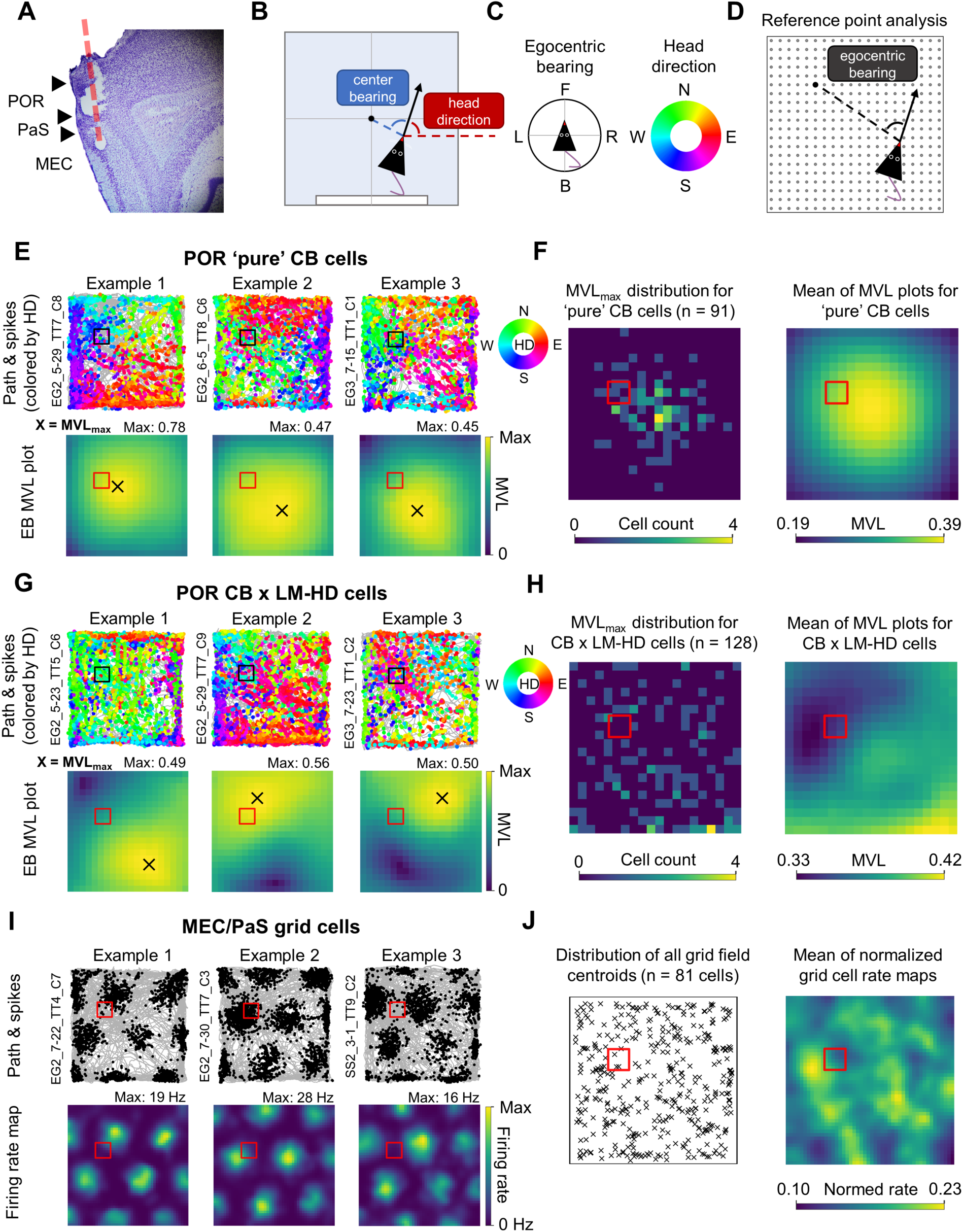
Unbiased spatial representation of task environment by POR and MEC/PaS neurons. **A)** Nissl-stained sagittal section from one rat (SS2) indicating the electrode path (red dotted line) as well as the estimated delineation between POR, PaS, and MEC (black triangles). **B)** Top-down schematic representation of the measurement of egocentric center-bearing, which is measured relative to the geometric centroid of the environment, and allocentric head direction, which is measured relative to the positive x-axis. **C)** Compass specifying the measurement of egocentric bearing (*left*; F = front, L = left, B = back, R = right) and compass/color code specifying the measurement of head direction (*right*; N = north, W = west, S = south, E = east). **D)** Schematic representation of the egocentric bearing reference point analysis used to compute egocentric tuning strength relative to different locations in the environment. Egocentric bearing was measured relative to a 20 x 20 array of evenly spaced potential reference points (gray dots). In the figure, egocentric bearing is being measured relative to a location in the northwest quadrant (indicated by a black dot). **E)** HD-colored path and spike plots (*top row*) and egocentric bearing mean vector length (MVL) plots (*bottom row*) for three example POR CB cells without conjunctive LM-HD tuning recorded during the standard navigation task. MVL plots indicate the strength of the cell’s tuning to the egocentric bearing of potential reference points spread across the environment, with the location of the highest value (MVL_max_ location) indicating the cell’s most strongly encoded reference point. **F)** *Left*, 2D histogram indicating the distribution of MVL_max_ locations for all ‘pure’ CB cells; *right*, mean of normalized MVL plots for all ‘pure’ CB cells. Note that MVL_max_ locations and overall MVL values are concentrated in the center of the environment. **G)** Same as **(E)** but for three POR CB cells with conjunctive LM-HD tuning. Note that their MVL_max_ locations are offset from the center of the environment. **H)** Same as **(F)** but for conjunctive CB x LM-HD cells. Note that MVL_max_ locations are scattered uniformly across the environment, with neither MVL_max_ locations nor overall MVL values showing bias toward the goal zone. **I)** Path and spike plots (*top row*) and 2D allocentric firing rate maps (*bottom row*) for three grid cells recorded from MEC/PaS during standard task sessions. **J)** *Left*, distribution of the centroids of all grid fields for all MEC/PaS grid cells recorded in the standard task; *right*, mean of normalized 2D allocentric firing rate maps for all grid cells. Note that neither grid field centroids nor overall grid cell firing rates showed clear bias toward the goal zone. Population distributions shown in **(F, H, J)** are based on resampled data that equalized occupancy across the environment (raw distributions shown in Fig. S3, S4, S5).

**Table 1.** Cell classifications by animal in the baseline recording session.

| **Animal** | **POR**<br>**total** | **‘Pure’**<br>**CB** | **CB x**<br>**LM-HD** | **CD** | **MEC/PaS**<br>**total** | **HD** | **Grid** | **Border** | **Non-grid**<br>**spatial** |
| --- | --- | --- | --- | --- | --- | --- | --- | --- | --- |
| SS1* | 95 | 18 | 32 | 22 | - | - | - | - | - |
| SS2 | 35 | 6 | 8 | 0 | 98 | 45 | 38 | 3 | 30 |
| EG2 | 99 | 28 | 37 | 18 | 137 | 86 | 29 | 6 | 49 |
| EG3 | 186 | 39 | 51 | 26 | 51 | 44 | 14 | 7 | 20 |
| **Total** | **415** | **91** | **128** | **66** | **286** | **175** | **81** | **16** | **99** |

\*Unit isolation for SS1 was lost before tetrodes crossed into MEC/PaS

We first investigated whether POR egocentric cells displayed tuning biases toward the learned goal location. While POR CB cells tend to bias their directional tuning toward the center of the environment (LaChance et al., 2019), we have proposed previously that combining CB and HD signals could allow for coding of specific allocentric locations by shifting the cell’s egocentric reference point from the environment center to some other location (LaChance and Taube, 2023a). If this were the case, we might expect POR cells conjunctively encoding CB and HD (CB x LM-HD cells) to show bias in their preferred reference locations toward the goal zone. To investigate this possibility, for each cell, we computed egocentric bearing tuning curves for a 20 x 20 array of potential egocentric bearing reference points spread evenly across the environment (Fig. 2D), and extracted the location with the highest mean vector length (MVL_max_ location; Wang et al., 2018) as the cell’s preferred egocentric reference point. As expected, ‘pure’ POR CB cells without conjunctive LM-HD tuning (n = 91) had MVL_max_ locations that clustered around the center of the environment (Fig. 2E, F; Fig. S3A). Critically, POR conjunctive CB x LM-HD cells (n = 128) displayed MVL_max_ locations that were offset from the environment center, but instead of clustering near the goal location, they were uniformly distributed across the environment (Fig. 2G, H; Fig. S3D). We also analyzed the spatial distribution of egocentric bearing MVLs across the environment for both CB and CB x LM-HD cells to assess if MVLs were generally higher near the goal zone. No MVL difference was observed between the goal zone and rotationally equivalent zones for ‘pure’ CB cells (Fig. 2F; Fig. S3B, C), but for CB x LM-HD cells the MVLs were found to be significantly lower in the goal zone than the equivalent zones (Fig. S3E, F). However, this effect seemed to be driven by the animals’ biased spatial occupancy near the goal zone, as it was no longer present after resampling the data to match occupancy time across the full environment (Fig. 2H; Fig. S3F).

To assess if overall firing rates of POR neurons showed bias toward the goal zone, we constructed normalized 2D allocentric firing rate maps for POR CB and CB x LM-HD cells and compared the firing rates in the goal zone vs. rotationally equivalent zones for all cells of each group. Neither group exhibited firing rate biases toward the goal zone (Fig. S4A-D). We also analyzed POR cells with significant tuning to the egocentric distance of the center/boundaries of the environment (center-distance (CD) cells; n = 66), which similarly did not exhibit any significant firing rate bias near the goal zone (Fig. S4E, F).

We next investigated whether MEC/PaS location- and HD-tuned cells displayed tuning biases toward the goal location. Grid cells have been previously shown to cluster their fields near dynamically shifting goals (Boccara et al., 2019), and both grid and non-grid spatial cells have been shown to display increased firing rates near an unmoving goal zone that also serves as a reward zone (Butler et al., 2019). To test for goal-related field concentration or increased firing rates in the current task, we 1) computed the locations of all firing field centroids for grid (n = 81), border (n = 16), and non-grid spatial (n = 99) cells and assessed concentration of field centroids near the goal zone vs. rotationally equivalent zones; and, 2) computed normalized allocentric firing rate maps for all grid, border, and non-grid spatial cells and compared their firing rates between the goal zone and equivalent zones. Neither method revealed biases toward the goal zone among grid cells (Fig. 2I; Fig. S5A-C), border cells (Fig. S5D-F), or non-grid spatial cells (Fig. S5G-I), suggesting that the MEC/PaS spatial cells uniformly represented the full task space rather than overrepresenting the goal location. Resampling the data to match spatial occupancy across the full environment (to control for behavioral bias toward the goal zone) produced the same pattern of results (Fig. 2J; Fig. S5).

Finally, we also assessed the distribution of preferred firing directions for POR LM-HD cells (n = 192) and MEC/PaS HD cells (n = 175), which also lacked bias toward the general direction of the goal zone (Fig. S6A, C). MVLs for HD-tuned cells were also not higher in the general direction of the goal zone (Fig. S6B, D), although POR LM-HD cells tended to have higher MVLs in the general direction of the cue card, consistent with previous reports of their visual landmark biases (LaChance et al., 2022). Overall, none of the analyzed POR nor MEC/PaS cell types exhibited spatial firing biases relative to the goal zone despite clear behavioral bias toward that location.

### Cortical dissociation of spatial reference frames during visual cue manipulation

Our results so far indicate a uniform spatial representation of the task environment in POR and MEC/PaS, despite strong behavioral bias toward a learned location. However, it is still unclear whether the POR and MEC/PaS spatial representations are based on the perceived configuration of local visual landmarks (i.e., relative to the perceived cue card location) or based on an internal model of location and orientation within the global environment (i.e., relative to the layout of the whole room). To differentiate these possibilities, we placed the visual scene in conflict with the room layout by changing the cue card location without moving the goal zone.

Each day began with a recording session using the standard cue configuration (standard session). This session was followed by a subsequent session in which the cue card was either moved to an adjacent wall (cue rotation sessions) or duplicated along the north wall (cue duplication session).

A final standard session then took place. Animals were always placed into the environment at the center of the south wall (facing north) to help them maintain a sense of global orientation despite visual cue changes. Accordingly, we found that animals were able to largely ignore the visual cue manipulations, continuing to approach the correct zone compared to geometrically equivalent zones in terms of both entries per minute and occupancy time following counterclockwise cue rotation (Fig. 3A-C), clockwise cue rotation (though the occupancy difference between the correct goal zone and the northeast equivalent zone was not significant (*P* = 0.06); Fig. 3D-F), and cue duplication (Fig. 3G-I). It is possible that the animals may have been slightly confused during the clockwise cue rotation session due to the cue card moving closer to and interfering with the goal zone, although their behavior was still generally directed toward the correct location.

**Figure 3.**
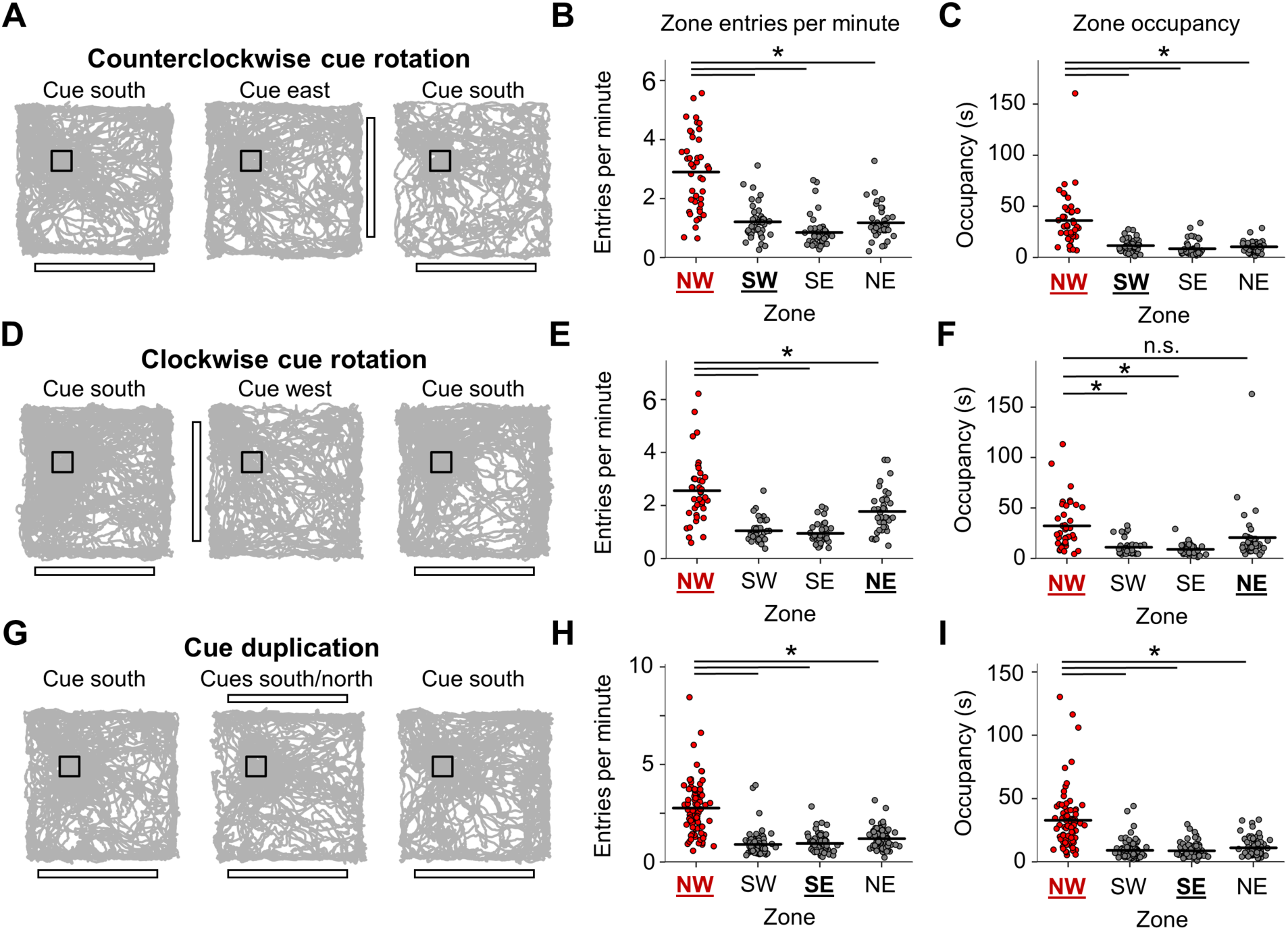
Consistent navigational performance despite visual cue manipulation. **A)** Example trajectory plots for one three-session round of the counterclockwise cue rotation experiment. **B)** Strip plot showing the number of entries per minute into the goal zone compared with rotationally equivalent zones. If animals followed the cue rotation, they would have entered more often into the southwest equivalent zone (bolded and underlined in black; repeated measures ANOVA, *F*(3, 123) = 68.12, *P* = 6.15e-17; paired *t*-tests, NW vs. SW, *t*(41) = 9.74, *P* = 9.61e-12; NW vs. SE, *t*(41) = 9.84, *P* = 7.05e-12; NW vs. NE, *t*(41) = 8.51, *P* = 4.00e-10). **C)** Same as **(B)** but for occupancy time in the goal zone compared to equivalent zones (repeated measures ANOVA, *F*(3, 123) = 37.33, *P* = 8.74e-9; paired *t*-tests, NW vs. SW, *t*(41) = 6.08, *P* = 9.85e-7; NW vs. SE, *t*(41) = 6.75, *P* = 1.11e-7; NW vs. NE, *t*(41) = 6.37, *P* = 3.83e-7). **D-F)** Same as **(A-C)** but for the clockwise cue rotation experiment (entries per minute: repeated measures ANOVA, *F*(3, 114) = 39.48, *P* = 1.30e-12; paired *t*-tests, NW vs. SW, *t*(38) = 7.62, *P* = 1.09e-8; NW vs. SE, *t*(38) = 7.95, *P* = 4.00e-9; NW vs. NE, *t*(38) = 3.78, *P* = 0.0016; occupancy: repeated measures ANOVA, *F*(3, 114) = 15.00, *P* = 2.12e-6; paired *t*-tests, NW vs. SW, *t*(38) = 5.58, *P* = 6.49e-6; NW vs. SE, *t*(38) = 5.99, *P* = 1.78e-6; NW vs. NE, *t*(38) = 2.41, *P* = 0.063). **G-I)** Same as **(A-C)** but for the cue duplication experiment (entries per minute: repeated measures ANOVA, *F*(3, 234) = 107.18, *P* = 2.70e-26; paired *t*-tests, NW vs. SW, *t*(78) = 13.02, *P* = 8.99e-21; NW vs. SE, *t*(78) = 11.42, *P* = 7.66e-18; NW vs. NE, *t*(78) = 10.17, *P* = 1.79e-15).

For POR cells recorded in the cue manipulation sessions, we focused on cells with LM-HD and conjunctive CB x LM-HD tuning, as POR LM-HD cells have been previously shown to lock their directional preferences to salient visual landmarks during random foraging (LaChance et al., 2022). If this were the case in the current experiment, we would expect POR LM-HD cells to rotate their preferred firing directions along with the cue card in the cue rotation session and become tuned to two opposite directions in the cue duplication session. This result is what we observed. POR LM-HD cells (including conjunctive cells) rotated their preferred firing directions along with the cue card in the cue rotation session (Fig 4A, B, E; Table 2) and became bidirectionally tuned in the cue duplication session compared to both standard sessions (Fig. 4G, H; Table 2), though bidirectionality remained slightly elevated in the second standard session compared to the initial standard session (Fig. 4H). While the rotation of POR LM-HD preferred directions did not statistically differ from the full 90° expected from the cue rotation (Fig. 4E), it should be noted that they numerically under-rotated on average (mean preferred HD shift, counterclockwise: 51.40 ± 39.25°; clockwise: -63.05 ± 37.04°). Likewise, in the cue duplication session, LM-HD cells were more strongly modulated by the original south cue than the new north cue (Fig. 4I), suggesting that they were not purely driven by local visual cues and may have incorporated some global directional information.

**Figure 4.**
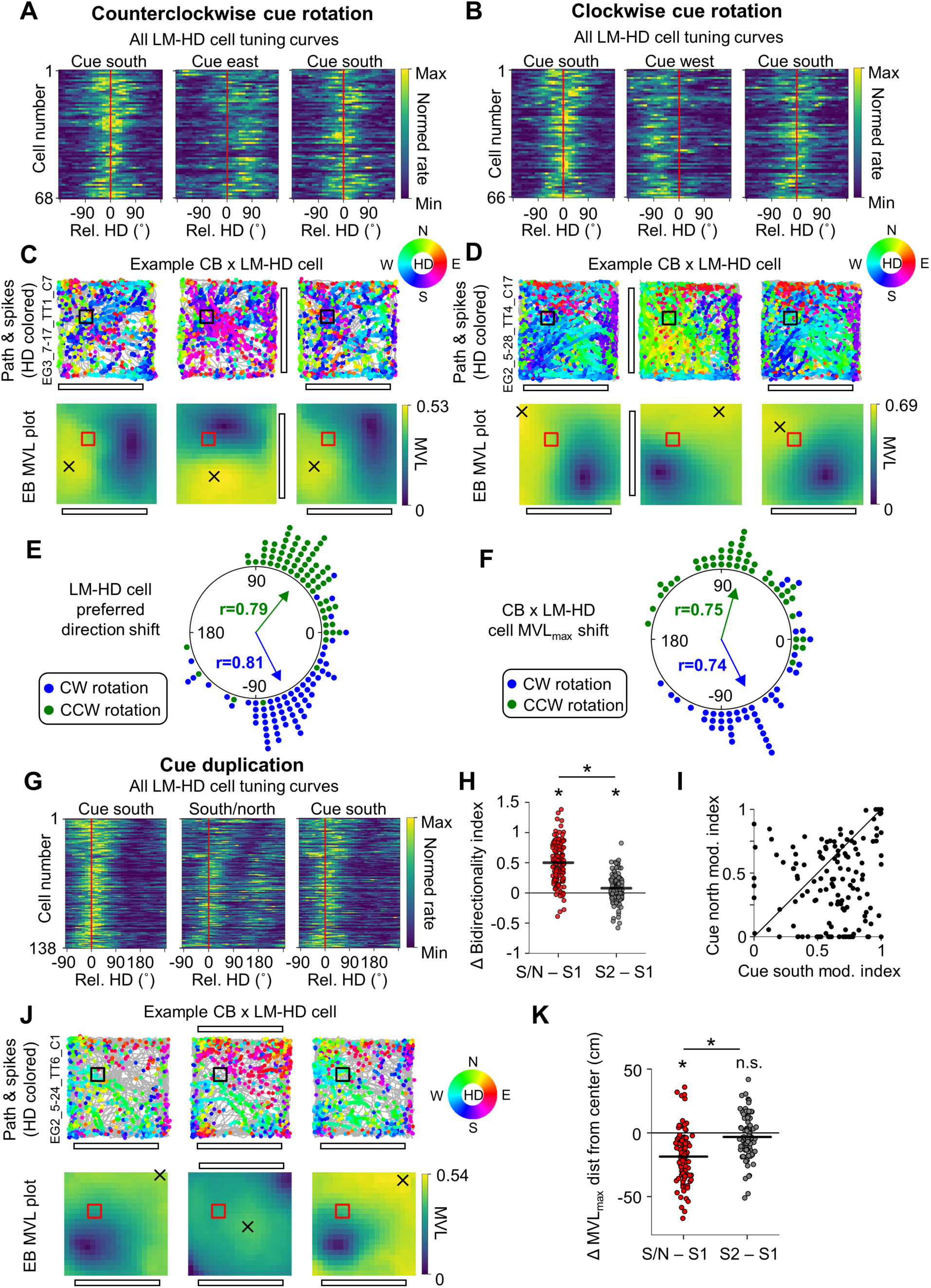
Postrhinal spatial representations follow visual cues despite unchanged behavior. **A)** Normalized tuning curves for all POR LM-HD cells recorded across all three sessions of the counterclockwise cue rotation experiment (n = 68 cells). Tuning curves have been realigned such that their mean direction is centered at 0, indicated by a red line. Note that the tuning curves shifted counterclockwise in alignment with the cue rotation. **B)** Same as **(A)** but for the clockwise rotation experiment (n = 66 cells). Note that HD tuning curves shifted clockwise in this condition. **C)** HD-colored path and spike plots (*top row*) and MVL plots (*bottom row*) for an example POR CB x LM-HD cell recorded in the counterclockwise rotation experiment. Note that the cell’s MVL_max_ location rotated around the center of the environment in alignment with the cue rotation. **D)** Same as **(C)** but for an CB x LM-HD cell recorded in the clockwise rotation experiment. Note that the MVL_max_ location rotated clockwise in this condition. **E)** Polar dot plot showing the change in preferred firing direction between the initial standard session and the following cue rotation session for all POR LM-HD cells. Note that the rotation of preferred directions did not statistically differ from the 90° visual cue rotation (counterclockwise: Rayleigh test, *r* = 0.79, *P* = 3.37e-19; V-test for concentration near 90°, *u* = 7.21, *P* = 2.84e-13; clockwise: Rayleigh test, *r* = 0.81, *P* = 1.34e-19; V-test for concentration near -90°, *u* = 8.31, *P* ≈0). **F)** Same as **(E)** but for the rotation of the MVL_max_ location around the center of the environment for POR CB x LM-HD cells. Note that the MVL_max_ rotation did not statistically differ from the 90° visual cue rotation (counterclockwise (n = 46 cells): Rayleigh test, *r* = 0.75, *P* = 1.40e-11; V-test for concentration near 90°, *u* = 6.91, *P* = 2.39e-12; clockwise (n = 45 cells): Rayleigh test, *r* = 0.74, *P* = 4.83e-11; V-test for concentration near -90°, *u* = 6.22, *P* = 2.53e-10). **G)** Same as **(A)** but for POR LM-HD cells recorded in the cue duplication experiment (n = 138 cells), and with preferred directions aligned toward the left side of the plot. Note that the tuning curves became visibly bidirectional when the cue was duplicated along the north wall. **H)** Change in bidirectionality index for all LM-HD cells between the initial standard session (S1) and both the cue duplication session (S/N) and the final standard session (S2). Note that cells became significantly more bidirectional in the duplication session compared to the standard sessions (repeated measures ANOVA, *F*(2, 274) = 212.47, *P* = 5.26e-46; paired *t*-tests, south/north vs. south 1, *t*(137) = 17.39, *P* = 5.23 e-36; south/north vs. south 2, *t*(137) = 14.31, *P* = 1.66e-28) and also remained slightly bidirectional in the second standard session (south 2 vs. south 1, *t*(137) = 4.18, *P* = 1.56e-4). **I)** Scatter plot showing the degree of firing rate modulation attributable to the south vs. north cues during the cue duplication session for all POR LM-HD cells. Note that most cells show some degree of modulation by both cues, but overall the population was more strongly modulated by the original (south) cue (paired *t*-test, *t*(137) = 6.55, *P* = 1.07e-9). **J)** Same as **(C)** but for a CB x LM-HD cell recorded in the cue duplication experiment. Note the bimodal distribution of HDs and the shift of the cell’s MVL_max_ location toward the center of the environment in the duplication session. **K)** Strip plot showing the change in distance of MVL_max_ locations from the center of the environment for all POR CB x LM-HD cells (n = 101 cells) between the initial standard session (S1) and both the cue duplication session (S/N) and the final standard session (S2). Note that MVL_max_ locations shifted significantly toward the center of the environment in the cue duplication session compared to both standard sessions (repeated measures ANOVA, *F*(2, 200) = 51.84, *P* = 5.22e-17; paired *t*-tests, south/north vs. south 1, *t*(100) = -9.31, *P* = 9.48e-15; south/north vs. south 2, *t*(100) = -6.97, *P* = 3.52e-10; south 2 vs. south 1, *t*(100) = -1.97, *P* = 0.15).

**Table 2.** Number of relevant spatial cells recorded during each cue manipulation.

| **Cue**<br>**manipulation** | **POR** |  |  | **MEC/PaS** |  |  |  |
| --- | --- | --- | --- | --- | --- | --- | --- |
|  | All<br>LM-HD | CB x<br>LM-HD | ‘Pure’<br>CB | HD | Grid | Border | Non-grid<br>spatial |
| CCW rotation | 68 | 46 | 30 | 57 | 41 | 3 | 28 |
| CW rotation | 66 | 45 | 26 | 66 | 24 | 7 | 37 |
| Duplication | 138 | 101 | 59 | 124 | 55 | 12 | 74 |

Critically, CB x LM-HD cells also displayed a rotation of their MVL_max_ locations around the center of the environment in response to cue rotation that did not differ statistically from 90° (Fig. 4C, D, F; Fig. S7; Table 2), though they also numerically under-rotated on average (mean MVL_max_ rotation, counterclockwise: 74.27 ± 43.60°; clockwise: -63.28 ± 45.09°). During the cue duplication session, MVL_max_ locations for CB x LM-HD cells shifted toward the environment center in response to the bipolar configuration of the visual cues (Fig. 4J, K; Table 2). Interestingly, ‘pure’ CB cells also exhibited a statistically significant rotation of their MVL_max_ locations in the cue rotation sessions (Fig. S8, S9A; Table 2), suggesting that they possessed minor visual landmark tuning properties that were undetected in the initial standard session. However, ‘pure’ CB cells did not show a significant shift of their MVL_max_ locations toward the environment center in the cue duplication session (Fig. S8, S9B; Table 2), likely due to the already close proximity of their MVL_max_ locations to the environment center (Fig. 2F).

In stark contrast with POR cells, the firing preferences of grid cells (Fig. S10), border cells (Fig. S11), non-grid spatial cells (Fig. S12), and HD cells in MEC/PaS (Table 2) were unchanged by the cue rotation or cue duplication manipulations, apparently maintaining a global sense of location and orientation within the overall room instead of following the configuration of visual cues. Grid, border, and non-grid spatial cell firing rate maps were highly correlated between the initial standard session, cue manipulation sessions, and the final standard session (Fig. 5C, D, F, J, K; Fig. S13). In addition, MEC/PaS HD cells tended to maintain their unidirectional firing preferences across sessions, failing to rotate in response to cue rotation (Fig. 5A, B, E). Interestingly, MEC/PaS HD cells displayed slightly elevated bidirectionality in both the cue duplication session and second standard session compared to the initial standard session, but there was no difference between the duplication session and second standard session (Fig. 5G-I), and the increase in bidirectionality in the duplication session was minor compared to that exhibited by POR LM-HD cells (mean ,1 bidirectionality index, MEC/PaS: 0.06 ± 0.21; POR: 0.50 ± 0.34). Thus, despite the animals displaying generally consistent behavior across all sessions of the cue manipulation experiment, altering the layout of local visual landmarks triggered a strong dissociation between POR cells that largely reflected the configuration of visual cues and MEC/PaS cells that largely reflected the animal’s ‘true’ global orientation relative to the goal zone and overall room.

**Figure 5.**
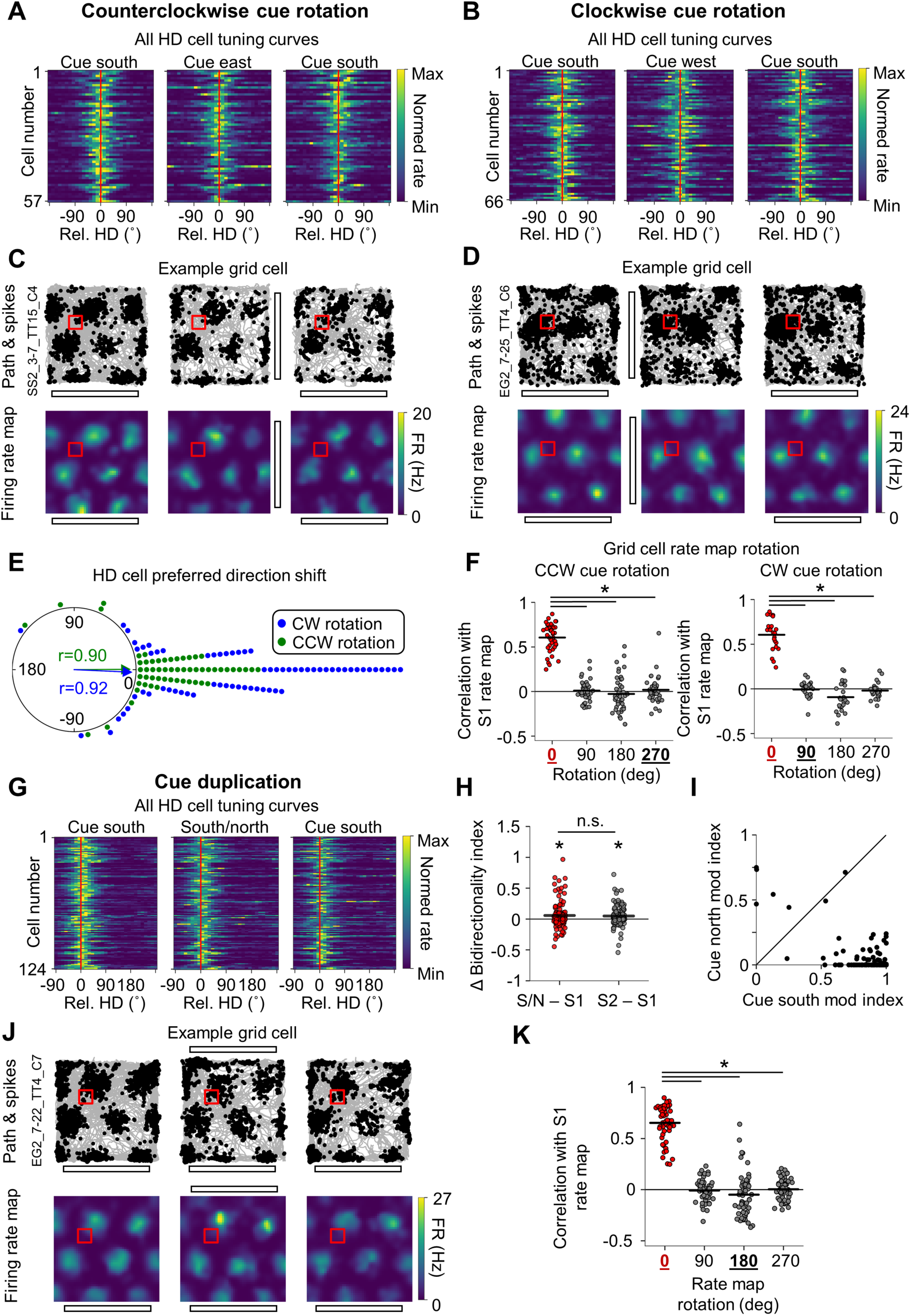
MEC/PaS spatial representations align with behavior despite visual cue manipulation. **A)** Normalized tuning curves for all MEC/PaS HD cells recorded across all three sessions of the counterclockwise cue rotation experiment (n = 57 cells). Tuning curves have been realigned such that their mean direction is centered at 0, indicated by a red line. Note that the were unchanged in response to cue rotation. **B)** Same as **(A)** but for the clockwise rotation experiment (n = 66 cells). Note that HD tuning curves were unchanged in this condition. **C)** Path and spike plots (*top row*) and 2D allocentric firing rate maps (*bottom row*) for an example MEC/PaS grid cell recorded in the counterclockwise rotation experiment. Note that the cell’s firing properties were unchanged in the rotation session. **D)** Same as **(C)** but for a grid cell recorded in the clockwise rotation experiment, also unchanged by the cue rotation. **E)** Polar dot plot showing the change in preferred firing direction between the initial standard session and the following cue rotation session for all MEC/PaS HD cells. Note that the preferred directions did not rotate in response to either direction of cue rotation. **F)** Strip plots for the counterclockwise experiment (*left*) and the clockwise experiment (*right*) showing correlations between the initial standard session rate map and the rotation session rate map that had been rotated in 90° increments, for all MEC/PaS grid cells. The rotation that would be expected to show high correlations if the rate maps rotated with the visual cue are bolded and underlined in black. Note that the grid cell rate maps did not rotate with the visual cue (counterclockwise (n = 41 cells): repeated measures ANOVA, *F*(3, 120) = 171.49, *P* = 6.42e-34; paired *t*-tests, 0° vs. 90°, *t*(40) = 19.31, *P* = 2.10e-21; 0° vs. 180°, *t*(40) = 14.90, *P* = 1.94e-17; 0° vs. 270°, *t*(40) = 19.93, *P* = 6.71e-22; clockwise (n = 24 cells): repeated measures ANOVA, *F*(3, 69) = 136.12, *P* = 9.64e-17; paired *t*-tests, 0° vs. 90°, *t*(23) = 19.77, *P* = 1.87e-15; 0° vs. 180°, *t*(23) = 11.84, *P* = 8.64e-11; 0° vs. 270°, *t*(23) = 16.24, *P* = 1.28e-13). **G)** Same as **(A)** but for MEC/PaS HD cells recorded in the cue duplication experiment (n = 124 cells), and with preferred directions aligned toward the left side of the plot. Note that the tuning curves were not visibly changed by cue duplication. **H)** Change in bidirectionality index for all HD cells between the initial standard session and both the cue duplication session and the final standard session. Note that bidirectionality was slightly increased in the duplication session and second standard session compared to the initial standard session (repeated measures ANOVA, *F*(2, 246) = 7.37, *P* = 0.0011; paired *t*-tests, south/north vs. south 1, *t*(123) = 3.09, *P* = 0.0073; south 2 vs. south 1, *t*(123) = 3.44, *P* = 0.0024) but did not differ between the duplication session and second standard session (south/north vs. south 2, *t*(123) = 0.47, *P* > 0.99). **I)** Scatter plot showing the degree of firing rate modulation attributable to the south vs. north cues during the cue duplication session for all MEC/PaS HD cells. Note that cells were much more strongly modulated by the south cue (paired *t*-test, *t*(123) = 27.44, *P* = 2.87e-54). **J)** Same as **(C)** but for a grid cell recorded in the cue duplication experiment. Note that cell firing was unchanged in the duplication session. **K)** Same as **(F)** but for the cue duplication experiment. Note that grid cell rate maps did not rotate in the cue duplication session (n = 55 cells; repeated measures ANOVA, *F*(3, 162) = 281.26, *P* = 1.97e-44; paired *t*-tests, 0° vs. 90°, *t*(54) = 28.88, *P* = 4.92e-32; 0° vs. 180°, *t*(54) = 18.82, *P* = 6.58e-25; 0° vs. 270°, *t*(54) = 24.34, *P* = 2.65e-30).

As MEC/PaS neurons were necessarily recorded after POR neurons due to the trajectory of the recording drive, we investigated whether the difference in visual cue responses could be attributed to different amounts of experience with the task rather than differences between the brain areas (i.e., perhaps POR representations rotated with the cue because the animal was less familiar with the task when the POR neurons were recorded). We used a linear mixed model to assess the relationship between task experience and the absolute shift of HD/LM-HD cell preferred firing directions following cue rotations (Fig. 6A, B) and between task experience and HD/LM-HD cell bidirectionality following cue duplication (Fig. 6C, D) while accounting for which brain area the HD or LM-HD cells were recorded from. No significant effect of the number of days of task experience was found for either rotation or duplication experiments, which, along with the abrupt change in response properties after crossing the ventral POR border (Fig. 6A, C), suggests that the difference in neural response properties was because the neurons were recorded from different brain areas instead of owing to different levels of experience with the task.

**Figure 6.**
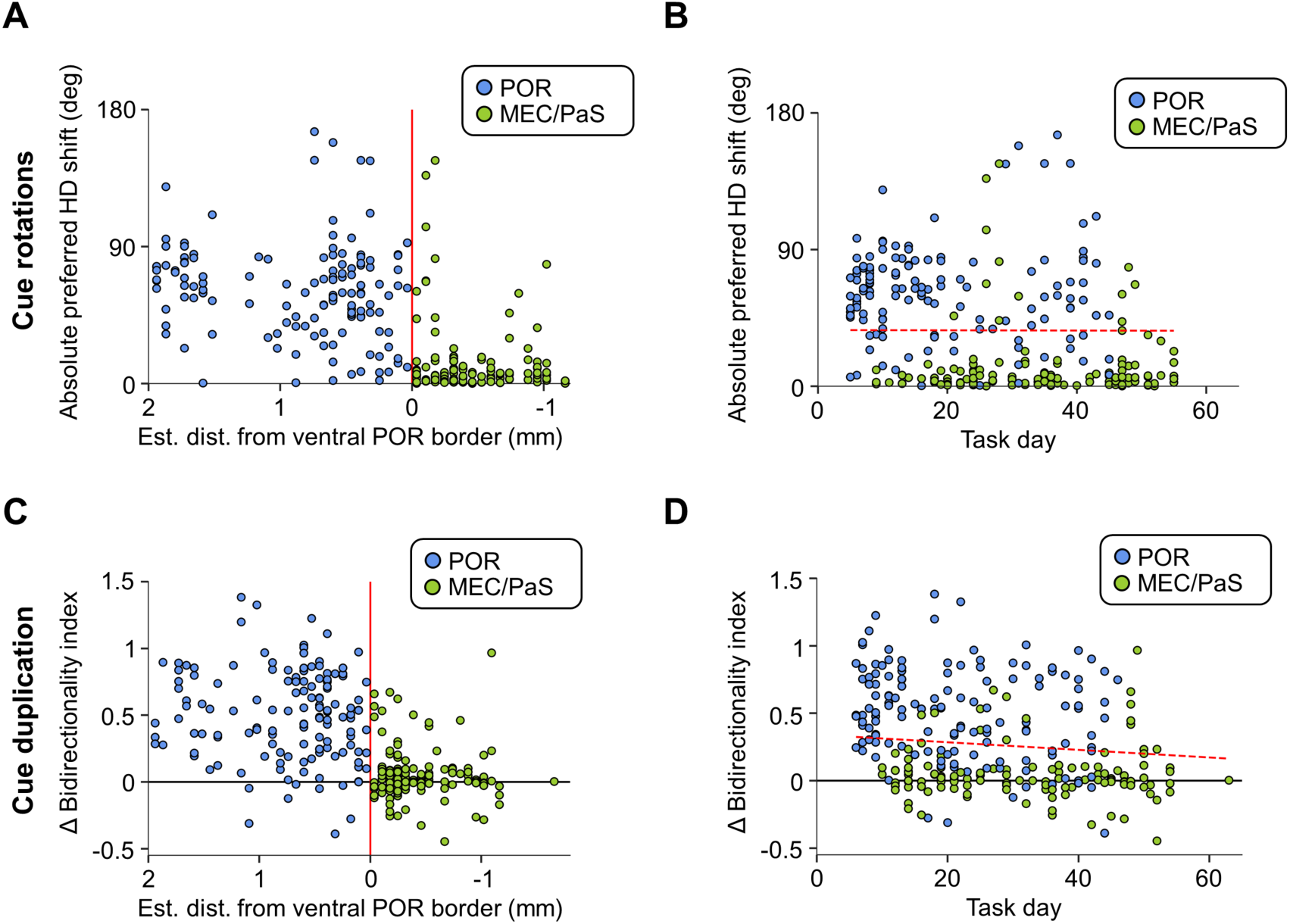
Abrupt change in visual cue responses at the ventral POR border. **A)** Scatter plot indicating the absolute shift in HD cell preferred directions during either clockwise or counterclockwise visual cue rotations compared to the estimated distance from the ventral POR border (indicated by a red line). Note the abrupt decrease in rotation extent as the tetrodes appeared to cross this border. **B)** Same as **(A)** but showing the absolute HD shift following cue rotation compared to the number of days the animals had engaged in the standard task. The red dotted line indicates the relationship between HD shift and task day based on the output of a linear mixed model that also considered brain area. No significant relationship was found between task day and shift magnitude (*Z* = -0.052, *P* = 0.96). **C-D)** Same as **(A-B)** but for change in bidirectionality index observed between the initial standard session and the following cue duplication session in the cue duplication experiment. Note the abrupt decrease in bidirectionality as the tetrodes appeared to cross the ventral POR border, and the lack of a significant correlation between bidirectionality and task day using a linear mixed model (*Z* = -0.044, *P* = 0.97).

## Discussion

This study investigated the neural encoding of spatial reference frames during a place navigation task in which entry into a learned, uncued goal zone in an open field environment triggered food reward to appear in a random location. While animal behavior was strongly biased toward the goal zone, spatial representations of single neurons in both POR and MEC/PaS encoded the task space uniformly. Subsequent rotation or duplication of a prominent local visual landmark without altering the goal zone did not affect behavioral bias toward the goal zone. Critically, however, POR egocentric spatial representations shifted to match the visual landmark configuration, while MEC/PaS allocentric spatial representations remained anchored to the room reference frame. These results indicate a clear dissociation between POR egocentric and MEC/PaS allocentric spatial reference frames despite strong behavioral evidence that the animal recognized its true allocentric position with respect to a remembered place.

### Relation to previous research on POR spatial representations

POR receives dense visual and visuospatial input (Burwell and Amaral, 1998b; Burwell, 2000; Tomás Pereira et al., 2016; Beltramo and Scanziani, 2019), particularly from the superior colliculus via the lateral posterior thalamus (Tomás Pereira et al., 2016; Beltramo and Scanziani, 2019). It is therefore unsurprising that POR neurons are strongly modulated by the spatial layout of visual landmarks, which has been suggested by lesion studies (Bucci and Burwell, 2004; Davies et al., 2007; Zhang et al., 2010) and demonstrated among single neurons during random foraging tasks with no explicit memory component (LaChance et al., 2022; LaChance and Hasselmo, 2024). However, a major advance of the current study is the observation that POR neurons shift their spatial representations along with visual landmarks despite the animal’s overt behavioral bias toward a memorized, unchanging allocentric goal location. It is important to note that, as with previous random foraging studies (LaChance et al., 2022; LaChance and Hasselmo, 2024), POR spatial representations did not fully follow the visual landmark, tending to under-rotate in response to cue rotation and showing greater modulation along their initial directional preference compared to their newly added preference in response to cue duplication. This incomplete response to visual landmark manipulation may be due to additional inputs to POR from parahippocampal and limbic areas carrying allocentric spatial signals (Burwell and Amaral, 1998b; Burwell, 2000; Agster and Burwell, 2013; Tomás Pereira et al., 2016) including the anterior thalamus, inactivation of which causes partial degradation of LM-HD (but not CB) tuning in POR (LaChance and Taube, 2024). However, it remains unclear if and how POR may contribute to place navigation given its overt encoding of task-irrelevant visual cues in the current study, especially as previous lesion studies have produced mixed results regarding the importance of POR in allocentric navigation tasks such as the Morris water maze (Liu and Bilkey, 2002; Burwell et al., 2004).

### Relation to previous research on MEC spatial representations

Initial investigations of spatially modulated cells in MEC, particularly grid cells, posited that they may form a regular metric for uniform mapping of the surrounding environment (Hafting et al., 2005; McNaughton et al., 2006; Moser et al., 2008). However, subsequent studies indicated that the typically uniform grid cell firing patterns could be distorted by manipulating behavioral affordances (Derdikman et al., 2009; Carpenter et al., 2015; Krupic et al., 2015; Krupic et al., 2018; Sanguinetti-Scheck and Brecht, 2020). Recently, studies employing different navigation tasks have demonstrated that grid and/or non-grid spatial signals in MEC can be influenced by nearby spatial goals (Boccara et al., 2019; Butler et al., 2019; Peng et al., 2023), a result that we did not observe in the current study. This distinction may be explained by differences in task design. In one study, grid fields were found to shift toward reward locations as animals learned to shuttle along stereotyped paths between three food wells that were newly selected each day (Boccara et al., 2019). In contrast, the current study had only one goal zone that was stable for weeks or months; goal-directed paths were highly variable across trials (discussed below); and the goal zone was not the same as the random reward location. Another study that was more similar to the current set of experiments found that grid and non-grid spatial cells exhibited higher firing rates near a single permanent goal zone that also served as the reward location (Butler et al., 2019). As the major difference between that study and the current study is that the reward location was random in the current study, the increased firing rates found near the goal zone in the previous study may be related to the animals’ anticipation of reward in that location. It has also been suggested that increased MEC firing near the goal zone in the previous study may constitute an ‘arrival at goal’ signal (Nyberg et al., 2022). However, as we did not observe this effect near the goal zone in our task, it is instead possible that the increased firing in the previous study constituted a ‘reward anticipation’ or ‘arrival at reward’ signal. Additionally, while a major result of this previous study was that location decoding based on MEC cell firing was more accurate near the goal location in the place navigation task compared to random foraging (Butler et al., 2019), we were unable to make the same comparison in our study due to the absence of a separate random foraging session in our paradigm. It is therefore possible that MEC/PaS spatial firing was subtly more accurate in the vicinity of the goal zone in the current study. Finally, a recent study recording grid cells during a path integration task found that grid cells anchored their firing patterns to a task-relevant landmark (Peng et al., 2023). This result is similar to our finding that grid cells remained anchored to the room reference frame (and therefore the goal zone) despite manipulation of task-irrelevant visual landmarks. It is possible that grid patterns in our task would have followed the visual landmarks if they predicted the location of the goal zone, which should be investigated in future studies. In any case, the fact that our MEC cells always remained aligned with the allocentric task structure is unsurprising given the importance of MEC for allocentric navigation tasks such as the Morris water maze (Nagahara et al., 1995; Burwell et al., 2004; Hales et al., 2014).

### Relation to hippocampal goal representations

Many previous studies of hippocampal place cells during real-world navigation tasks have reported overrepresentation of goal locations by place fields in both CA1 (Breese et al., 1989; Kobayashi et al., 1997; Hollup et al., 2001; Fyhn et al., 2002; Lee et al., 2006; Dupret et al., 2010; Danielson et al., 2016; Mamad et al., 2017; Zaremba et al., 2017; Turi et al., 2019; Xu et al., 2019; Kaufman et al., 2020; Xiao et al., 2020) and CA3 (Kobayashi et al., 2003; but see Dupret et al., 2010; for review, see O’Keefe and Krupic, 2021; Sosa and Giocomo, 2021; Nyberg et al., 2022). However, unlike the current study, these experiments generally required the animals to take repeated stereotyped goal-directed paths (Nyberg et al., 2022). In contrast, in studies where the animals needed to constantly plan novel goal-directed paths from a variety of locations (as with our task), overrepresentation of the goal zone has not been observed (Jeffery et al., 2003; Hok et al., 2007b; Pfeiffer and Foster, 2013; Spiers et al., 2018; Duvelle et al., 2019). These include studies with one or two long-term goal locations (as in the current study; Jeffery et al., 2003; Hok et al., 2007b; Duvelle et al., 2019) or with variable goal locations (Pfeiffer and Foster, 2013; Spiers et al., 2018). Therefore, it is not necessarily surprising that MEC cells, which provide dense input to the hippocampus (Steward and Scoville, 1976; Witter et al., 2000), and PaS cells, which have allocentric spatial properties and project strongly to MEC (van Groen and Wyss, 1990; Boccara et al., 2010), also failed to show overrepresentation of the goal zone in the current study. Despite the absence of place field overrepresentation in these types of tasks, some studies have found evidence for increased out-of-field firing by place cells in the goal zone when the animal was required to wait for 2 seconds in the zone before retrieving randomly scattered reward (Hok et al., 2007b; Hok et al., 2007a; Duvelle et al., 2019). As our task did not require the animals to wait in the goal zone to be rewarded, we could not investigate the possibility of increased out-of-field firing in MEC/PaS grid or non-grid spatial cells. However, future studies should investigate this possibility.

Aside from explicit place field overrepresentation, goal-direction representations have also been observed in the hippocampus (Sarel et al., 2017; Ormond and O’Keefe, 2022). However, given our observations that POR egocentric directional representations are primarily tied to task-irrelevant visual cues, in addition to POR only providing sparse direct innervation to the hippocampus (Naber et al., 2001), it seems unlikely that POR is a major driver of these hippocampal goal-direction representations.

### Relation to previous goal-directed navigation models

Early computational models of the neural mechanisms underlying place navigation focused on computations occurring within the hippocampus, such as route encoding by CA3 recurrent collaterals (Redish and Touretzky, 1998) or rotation of incoming egocentric maps using an incoming HD signal (Recce and Harris, 1996). While these models (sometimes implicitly) considered inputs from higher level sensory areas (e.g., POR) or the entorhinal cortex, they did not depend on goal-related information being contained in these inputs, which would instead be ‘purely’ sensory or ‘purely’ spatial, respectively. These models therefore agree with the absence of clear goal zone biases exhibited by the locational or directional firing of POR and MEC/PaS neurons in the current study. More recent models of entorhinal function during goal-directed navigation, which have largely focused on grid cell coding (Erdem and Hasselmo, 2012; Kubie and Fenton, 2012; Erdem and Hasselmo, 2014; Bush et al., 2015; Banino et al., 2018; Edvardsen et al., 2020), have also largely assumed that grid cell firing fields would not explicitly overrepresent goal locations, but rather that higher-level properties of grid cell populations, such as forward trajectory planning (Erdem and Hasselmo, 2012; Kubie and Fenton, 2012) that may relate to recently discovered entorhinal-hippocampal sweeps (Vollan et al., 2025), could be read out by downstream brain regions to drive or support navigation for both goal-finding and obstacle avoidance. Regarding POR, one previous model of goal-directed navigation (LaChance and Taube, 2023a) also assumed that it contained ‘purely’ sensory information that would combine with reward information downstream, potentially in the dorsal striatum. The results of our study are therefore largely aligned with predictions from several previous models of neurally driven goal-directed behavior, specifically that POR firing reflects egocentric information derived from sensory cues and MEC/PaS firing reflects the allocentric spatial layout of the environment.

### Implications for POR-MEC interactions

POR sends direct projections to MEC (Naber et al., 1997; Burwell and Amaral, 1998a; Witter et al., 2000; Koganezawa et al., 2015) and has been considered a major source of sensory input to MEC (Burwell and Amaral, 1998a; Eichenbaum et al., 2007; Knierim et al., 2014). It would therefore be reasonable to expect that MEC spatial representations would reflect POR inputs. In contrast, our study demonstrates that POR and MEC process the task environment via distinct reference frames, with POR largely reflecting the layout of nearby visual landmarks and MEC largely reflecting the animal’s global orientation relative to the goal zone and overall room. It remains unclear how POR representations actually impact downstream MEC neurons given this distinction. One possibility is that the ‘pure’ CB and CD cells, which seem to indicate the geometric structure of the environment rather than the visual landmark layout (LaChance and Taube, 2023b), could still support global allocentric coding in MEC if combined with a ‘classic’ HD signal originating elsewhere (e.g., from the postsubiculum (Taube et al., 1990) or the anterior thalamus (Taube, 1995; Winter et al., 2015)), though this would require MEC to somehow ignore the LM-HD signal originating in POR while still considering the CB/CD signal. Some models have suggested how MEC could use ‘classic’ HD input combined with egocentric coding of environmental boundaries to support allocentric spatial firing (Bicanski and Burgess, 2018; Alexander et al., 2023). However, we have previously demonstrated that ‘pure’ POR CB and CD cells alter their firing patterns in response to local changes in environmental geometry while grid cell firing is unaffected (LaChance and Hasselmo, 2024), making it unlikely that MEC firing properties are derived from simple operations (e.g., linear summation) performed on afferent egocentric boundary signals.

It is worth noting that POR neurons are not monolithic and have shown heterogeneity in their encoding of local vs. global spatial cues in prior studies (LaChance and Taube, 2023b; LaChance and Hasselmo, 2024). One possibility is that POR, which also projects to the lateral entorhinal cortex (LEC) in addition to MEC (Burwell and Amaral, 1998a; Doan et al., 2019), could preferentially contribute information about global cues to MEC while separately contributing information about local cues to the lateral entorhinal cortex (Neunuebel et al., 2013; Knierim et al., 2014; Wang et al., 2018). While distal visual cues in the current study were limited by a black curtain surrounding the arena, POR neurons may have exhibited stronger global firing properties if distal visual cues were available. Future studies should investigate the specific types of spatial information in POR that contribute to distinct spatial representations in each entorhinal subdivision, as well as the neural mechanisms underlying the proposed transformation from largely egocentric firing in POR to largely allocentric firing in MEC.

## Conclusion

Overall, the results of this study indicate an absence of spatial firing biases among POR and MEC/PaS cells during navigation relative to a well-learned, uncued allocentric place. They also reveal a strong dissociation between POR and MEC/PaS in their encoding of local and global spatial reference frames, respectively, when those reference frames are placed into conflict via visual landmark manipulation, such that POR neurons shift with local visual landmarks while MEC/PaS neurons remain locked to the globally defined allocentric goal location. Future studies should seek to illuminate the specific mechanisms underlying interactions between these regions as well as their distinct contributions to place learning.

## Acknowledgements

We thank Éléonore Duvelle for helpful comments on the manuscript. This work was supported by the National Institute of Mental Health, grant numbers F32 MH139270 (P.A.L.) and R01 MH120073 (M.E.H.).

## Data and code availability

Data and code needed to reproduce the findings of this paper will be uploaded to an online repository upon publication.

## Author Contributions

P.A.L. designed the experiments. P.A.L., S.S., and E.G. ran the experiments. P.A.L. performed the analyses and drafted the manuscript. P.A.L. and M.E.H. discussed and finalized the manuscript.

## Supplementary Materials

Figures S1-14

## Materials and Methods

### Subjects

Subjects were four female Long-Evans rats (Charles River Laboratories) aged 5-14 months and weighing 300-350 grams prior to surgery. Rats were individually housed in Plexiglas cages and maintained on a 12 h light/dark cycle. Before surgery, food and water were provided ad libitum. All experimental procedures were performed in compliance with institutional standards as set forth by the National Institutes of Health *Guide for the Care and Use of Laboratory Animals* and approved by the Boston University Institutional Animal Care and Use Committee.

### Electrode construction

Recording drives consisted of a moveable bundle of eight tetrodes. Tetrodes were constructed by twisting together four pieces of 17-μm nichrome wire (Alleima), which were then threaded through one 26-gauge piece of polyimide tubing affixed to the shuttle of a 3D-printed microdrive (Vöröslakos et al., 2021; print designs from https://github.com/buzsakilab/3d_print_designs) which could be advanced in the dorsal-ventral plane by turning a single 00-90 screw. The end of each wire was connected to one contact of a 32-channel electrode interface board (EIB; Neuralynx).

### Electrode implantation and recovery

Animals were first anesthetized with isoflurane. They were then placed into a stereotaxic frame, and an incision was made in the scalp to expose the skull. Anchor screws were fixed to the skull and secured with a layer of metabond. One anchor screw placed above the frontal cortex was used as a ground screw. A craniotomy was drilled above the postrhinal cortex (1.9 mm posterior and 4.6 mm lateral to lambda) to expose the anterior edge of the transverse sinus. The tetrode bundle was subsequently implanted 4.6 mm lateral to lambda, 0.40 mm anterior to the anterior edge of the transverse sinus, and ∼0.5 mm ventral to the cortical surface. The bundle was given a 10° angle in the anterior-posterior plane, such that the tetrode tips pointed forward. The drive body was secured to the skull using dental acrylic, and was surrounded by a 3D-printed headcap (Vöröslakos et al., 2021) which housed the EIB and was also secured to the skull using dental acrylic. Two animals (SS1 and SS2) received an additional tetrode bundle implant targeting the retrosplenial cortex; however, due to low cell yields, the associated data is not presented here. Animals were given 7 days to recover from surgery, after which they were placed on food restriction such that their body weight reached 85-90% of their pre-surgical weight.

### Place navigation task

The place navigation task took place in a 120 x 120 cm box with a black floor and 50 cm high black walls that was surrounded on all sides by a floor-to-ceiling circular black curtain. The box itself was featureless aside from a single white cardboard sheet (‘cue card’) placed along the center of the south wall. The cue card was 50 cm in height and 72 cm in width, such that it covered ∼60% of the horizontal extent of the wall. An overhead color video camera was used to track the animal’s location and orientation within the box based on the placement of red and green light-emitting diodes (LEDs) mounted on the animal’s head and spaced approximately 6 cm apart over the head and back of the animal, respectively. The locations of the two LEDs were monitored by Cheetah data acquisition software (Neuralynx) which also processed and saved the neural data (described below). We used Zoned Video software (Neuralynx) to define a “goal zone” within the arena, which was approximately 15 cm square and centered 35 cm from the west wall and 45 cm from the north wall for all animals. Zoned Video was used to detect entries into and exits out of the goal zone. Detection of an entry into the goal zone prompted Cheetah to send a TTL pulse to an Arduino microcontroller which controlled a sucrose pellet dispenser mounted approximately 3 m directly above the arena, prompting the release of 1-4 chocolate sucrose pellets (Envigo, 45mg; number of pellets selected randomly on each trial) which randomly scattered across the arena floor. Once reward release had been triggered, a timeout period of 10 seconds began which prevented zone entry from triggering reward until the timeout period was over, to discourage animals from repeatedly entering the goal zone in quick succession.

Training in the place navigation task began once animals had recovered from surgery and had been placed on food restriction. Initial training sessions involved allowing each animal to freely explore the task environment for periods of 40-60 minutes per day, which was decreased over time to 20 minutes as the animals became familiar with the task structure and showed evidence of repeatedly visiting the goal zone to trigger reward release. All task sessions following this initial training period were 20 minutes in length. Recording and analysis of neural data began once an animal was triggering reward release approximately once per minute (20 times per 20 minute session).

### Cue manipulation sessions

Once animals were receiving approximately one reward per minute in the standard place navigation task, we began changing the visual cue configuration without changing the goal zone in order to place the local visual scene in conflict with the global layout of the task space. These manipulations took one of three forms:

> *Counterclockwise cue rotation*: Following an initial 20 minute standard session with the cue card along the south wall, the cue card was moved to the east wall, and another 20 minute session was run. A final 20 minute standard session was then run with the cue along the south wall.
>
> *Clockwise cue rotation*: Following an initial 20 minute standard session with the cue card along the south wall, the cue card was moved to the west wall, and another 20 minute session was run. A final 20 minute standard session was then run with the cue along the south wall.
>
> *Cue duplication*: Following an initial 20 minute standard session with the cue card along the south wall, a second identical cue card was placed along the north wall, and another 20 minute session was run. A final 20 minute standard session was then run with the north cue removed.

Between sessions, animals were placed into a cardboard box outside the task environment while the cues were manipulated. The arena floor was wiped down with a veterinary-safe disinfectant spray (REScue) and allowed to dry fully before the start of the following session. Animals were always placed back into the arena in the same location and orientation (middle of the south wall facing north) to help them maintain a sense of their orientation within the overall room despite the changing visual scene inside the task environment.

### Recording of neural data

While the animals completed the place navigation task, over the course of weeks or months, tetrodes were ‘screened’ for units that displayed well-isolated waveforms. Electrical signals were pre-amplified using unit-gain operational amplifiers on an HS-36-LED headstage and sent to a Digital Lynx SX acquisition system (Neuralynx). Signals from each tetrode wire were differentially referenced to a relatively quiet, low noise channel from a separate tetrode and bandpass filtered (600 Hz to 6 kHz) using Cheetah data acquisition software (Neuralynx). If waveforms on a given tetrode crossed a pre-defined amplitude threshold (typically 30 to 50 μV) they were timestamped and digitized at 32 kHz for 1 ms. A color video camera mounted above the arena captured video frames with a sampling rate of 30 Hz, which were timestamped so they could be matched to both the neural data and the timestamps of TTL pulses that signified entry or egress from the goal zone (described above). Videos were further analyzed using DeepLabCut (Mathis et al., 2018) to obtain a more accurate estimate of the LED positions offline. At least one 20 min recording session took place each day as the animal engaged in the place navigation task. Tetrodes were typically kept in place for at least two days so that the same group of cells could be recorded during both cue rotation and cue duplication manipulations (described below), after which they were advanced ∼50 to 100 μm and screened again the following day.

### Spike sorting

Spike sorting was conducted offline, with clustering initially performed by the automated clustering program Kilosort (Pachitariu et al., 2023). When cells were recorded across multiple sessions in a day, the sessions were first merged before being clustered by Kilosort to ensure cluster continuity, after which the sessions were separated for manual curation and analysis. Manual curation, which was not always necessary, involved visualizing waveform features including peak, valley, height, width, and principal components across multiple tetrode wires as a 3D scatter plot using the program SpikeSort3D (Neuralynx). Adjustment of automatically sorted clusters involved either merging clusters or drawing a polygon around the visually apparent boundaries of each cluster to exclude outlying spikes. Single-unit isolation was assessed using metrics such as L-ratio and isolation distance (Schmitzer-Torbert et al., 2005), as well as ensuring that temporal autocorrelograms contained a clear refractory period. As long as tetrodes were advanced between recording sessions, cells recorded across days were treated as independent units. Otherwise, the recording session with the larger number of isolated cells was used for baseline cell analyses.

### Histology

Following the completion of recordings, animals were deeply anesthetized with sodium pentobarbital, and small marking lesions were made at the tips of the tetrodes by passing a small anodal current (20 μA, ∼5 s) through each tetrode wire. Animals were then intracardially perfused with saline followed by 10% formalin solution, after which brains were removed from the skull and postfixed in 10% formalin for at least 24 hours. The brains were later transferred to 30% sucrose solution for at least 24 hours, after which they were frozen and sliced into 40 μm sagittal sections using a cryostat, with sections mounted onto glass microscope slides and subsequently Nissl-stained with cresyl violet. The location of the tetrode bundle over the course of the experiment was estimated by measuring backward from the most ventral portion of the marking lesions, which always ended in MEC or PaS (Fig. S14). Delineations of parahippocampal regions were drawn mainly from (Burwell, 2001; Boccara et al., 2010; Boccara et al., 2019). Due to minor spread of the tetrode tips and the variable slope of the ventral POR border, in some cases we judged individual tetrodes to have crossed the ventral POR border before others based on the sudden appearance of strongly theta-modulated cells and grid cells, which have been observed to exist in MEC/PaS but not POR (Hafting et al., 2005; Boccara et al., 2010; LaChance et al., 2019; LaChance and Hasselmo, 2024; Fig. S2). Note that one animal’s recording drive stopped acquiring well-isolated waveforms before the tetrodes exited POR (SS1; Fig. S14), so MEC/PaS cells were only recorded from three of the animals (Table 1).

### Initial cell classifications with a generalized linear model

We used 10-fold cross-validation with a Poisson generalized linear model (Hardcastle et al., 2017; LaChance et al., 2019) to classify individual cells as significantly encoding one of the five following variables: 2D allocentric location, allocentric head direction, egocentric bearing of the environment center (proxy for boundaries), egocentric distance of the environment center, and linear speed. the firing rate vector *r* for one cell across the full recording session was modeled as follows:

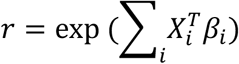

where *X* is a matrix containing animal state vectors for a single behavioral variable over time points *T, β* represents the parameter vector for that behavioral variable (similar to a tuning curve), and *i* indexes across behavioral variables included in the model. The parameter vectors were optimized by maximizing the log-likelihood *l* of the real spike train *n* given the estimated rate vector *r* across time points *t*:

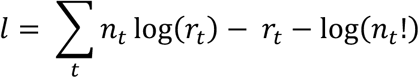

A small smoothing penalty, *P*, was added to the objective function to avoid artifacts and overfitting, which penalized differences between adjacent bins of each parameter vector:

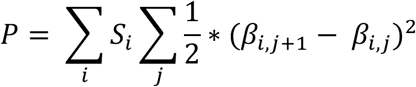

Here, *S* is a smoothing hyperparameter (set to 2 for 2D location and 20 for all other variables), *i* indexes over variables, and *j* indexes over response parameters for a given variable. Response parameters were estimated by minimizing (P – *l*) using SciPy’s *optimize.minimize* function. 400 bins (20 x 20 square) were used for the 2D location parameter vector, thirty bins were used for center bearing and allocentric head direction parameter vectors, and ten bins were used for center distance and linear speed.

Data for a session was split into training (9/10 of the session) and test (1/10 of the session) data (*k* = 10 folds). Parameter vectors were computed by minimizing the objective function on the training data using the full model with all four variables, to reduce potential correlative artifacts between independent variables (Burgess et al., 2005). Log-likelihood values for all possible variable combinations were computed. This procedure was repeated until all parts of the data had been used as test data.

Model selection followed a forward selection procedure (Hardcastle et al., 2017). Briefly, the log-likelihood values from the best two-variable model were compared to those from the best one-variable model. If the two-variable model showed significant improvement from the one-variable model (using a one-sided Wilcoxon signed-rank test), then the best three-variable model was compared to the two-variable model, and so on. If the more complex model was not significantly better, the simpler model was chosen. If the chosen model performed significantly better than a model that only included the cell’s mean firing rate, the chosen model was used as the cell’s classification. Otherwise, the cell was marked ‘unclassified’.

### Allocentric location firing rate maps

The animal’s two-dimensional location throughout the recording session was divided into 2.5 cm x 2.5 cm bins. For each cell, the number of spikes associated with each bin was divided by the amount of time the bin was occupied. The resulting firing rate map was smoothed with a Gaussian filter with a standard deviation of 3.75 cm.

## Classification of grid cells

After computing a cell’s allocentric firing rate map, it was used to compute a grid score (Hafting et al., 2005). A 2-dimensional autocorrelation was computed by correlating a copy of the rate map with the original rate map at all possible spatial shifts. A cell with hexagonally periodic firing fields would be expected to show a ring around the center of the autocorrelation with six evenly spaced peaks. For each cell, we identified the most probable inner and outer radii of this ring, and then computed a 1-dimensional autocorrelation by correlating a copy of the ring with the original ring at 3° rotational offsets from 0° to 180°. A grid score was then computed from the 1D autocorrelation by taking the lowest correlation value at 60° or 120° and subtracting the highest correlation value at 30°, 90°, or 150°. Cells that passed the GLM classification procedure for 2D location modulation and which had grid scores > 0.4 were considered grid cells.

### Classification of border cells

After computing a cell’s allocentric firing rate map, it was used to calculate a border score (Solstad et al., 2008). Border cells can also be considered a special case of boundary vector cells that exhibit a preference for proximal boundary distances (Barry et al., 2006; Lever et al., 2009). We first thresholded the rate map to exclude all bins with firing rates < 20% of the maximum firing rate. We then identified firing fields as groups of contiguous bins with an area of at least 200 cm^2^, after which we computed the maximum coverage of any single wall by any single firing field (expressed as a fraction *c*). We then computed the mean distance from the nearest wall of all bins belonging to any firing field (normalized by their firing rates), and normalized the resulting value by one half the longest length of any wall to calculate the value *d*. The border score was then computed as follows:

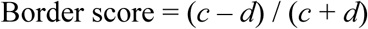

A neuron was considered a border cell if it passed the GLM classification procedure for 2D location modulation and had border score > 0.5.

### Classification of non-grid spatial cells

To classify cells without explicit grid or border tuning as having significant allocentric location tuning (Diehl et al., 2017), we computed the spatial information content (in bits/spike) for each cell. This was computed based on smoothed allocentric firing rate maps and smoothed spatial occupancy histograms using the following equation (Skaggs et al., 1997):

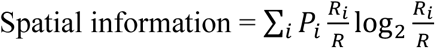

Where *i* indexes across spatial bins, *P_i_* is the probability of the animal occupying bin *i* (taken from the smoothed occupancy histogram), *R_i_* is the mean firing rate of the cell in bin *i* (taken from the smoothed firing rate map), and *R* is the mean firing rate across all spatial bins. A cell was classified as a non-grid spatial cell if it passed the GLM classification procedure for 2D location modulation, had spatial information content > 99^th^ percentile of a within-cell shuffle distribution (discussed below), and failed to be classified as a grid or border cell.

### Classification of head direction and landmark modulated-head direction cells

We computed a head direction tuning curve for each cell using 12° bins. The number of spikes fired by the cell in each bin was divided by the amount of the time that bin was occupied. We then computed the mean vector length of the resulting tuning curve to indicate tuning strength, while the mean angle indicated the cell’s preferred firing direction. A cell was classified as a head direction cell if it passed the GLM classification procedure for head direction modulation, had a mean vector length > 99^th^ percentile of a within-cell shuffle distribution as well as an overall threshold of 0.2, and had a peak firing rate > 1 Hz in its head direction tuning curve.

### Classification of center-bearing cells

Center-bearing tuning curves and cell classifications used the same criteria as for head direction cells, but based on the egocentric bearing of the environment center (i.e., the angle between the animal’s heading and the center of the environment). Tuning to the center can be considered a proxy for tuning to the boundaries of the environment.

### Classification of center-distance cells

To classify cells as being tuned to the distance of the center or boundaries of the environment, we computed the animal’s distance from the center of the environment and created a tuning curve using 4cm bins. The number of spikes fired by the cell in each bin was divided by the amount of the time that bin was occupied. We fit both linear and Gaussian functions to the tuning curve, and computed the R^2^ value of each fit to determine tuning strength. A cell was classified as center-distance cell if it passed the GLM classification procedure for egocentric distance tuning, had a linear *or* Gaussian R^2^ > 99^th^ percentile of a within-cell shuffle distribution, and had a peak firing rate > 1 Hz in its egocentric distance tuning curve.

### Egocentric mean vector length analyses

For each cell, we assessed egocentric bearing tuning relative to a 20 x 20 array of potential reference points spread evenly across the environment. Egocentric bearing tuning curves were created for each reference point, indicating the cell’s firing rate when that reference point occupied different angles relative to the animal’s heading. We computed the mean vector length of each tuning curve to indicate the tuning strength of the cell’s firing relative to each potential reference point, which could be visualized as a heat map (“MVL plot”). The location with the highest mean vector length was considered the cell’s ‘preferred’ reference location (“MVL_max_ location”). We also assessed overall egocentric bearing tuning strength relative to different reference points for a given group of cells by computing the mean of the MVL plots across the group of cells.

### Firing field centroid analyses

To assess the distribution of firing fields for grid, border, and non-grid spatial cells, we first computed a 2D allocentric rate map for each cell and subsequently thresholded the rate map to remove bins with firing rates < 30% of the maximum. We then identified firing fields as contiguous groups of bins with an area of at least 150 cm^2^, and computed the centroid of each firing field. The field centroids for each spatial cell type (grid, border, and non-grid spatial) were then visualized as a scatter plot.

### Spatial firing rate analyses

To assess the distribution of firing rates across the task space for different cell types, we first created 2D allocentric rate maps for each cell of a given type, and subsequently rescaled the rate maps to range from 0 to 1. We then computed the mean of the rate maps for the given cell type. For POR center-distance cells, which tend to either increase or decrease their firing rates with distance from the center of the environment in similar proportions (LaChance et al., 2019), we inverted the rate maps of cells with negative center-distance slopes before computing the overall mean.

### Spatial resampling analyses

As the animals spent the majority of each recording session near the goal zone, we sought to control for potential artifacts of this spatial bias when assessing neural tuning to egocentric bearing or spatial location. For example, occupying a specific location makes it difficult to accurately characterize the egocentric bearing of that location (i.e., how do you determine the angle between your heading and your current location?) which can lead to ‘flattening’ of the tuning curves constructed relative to that location. To correct for this issue, for a given recording session, we first separated the arena into 15 x 15 cm square bins (64 bins total) and identified the bin with the lowest occupancy time. We then resampled (without replacement) the tracking and spike data for all 64 bins, such that all bins had the same occupancy time. We then performed the egocentric bearing and/or spatial firing analyses on the resampled data.

### Assessment of HD/LM-HD cell bidirectionality

To assess if HD/LM-HD cells fired in two opposite directions in the cue duplication experiment, we computed a bidirectionality index (discussed in detail in (LaChance et al., 2022)). Briefly, two tuning curves were constructed for each HD/LM-HD cell: one based on the animal’s actual HD; and one where the animal’s HD had been doubled first. Symmetrical bidirectional distributions can be transformed into a unidirectional distribution by doubling the associated angles. The bidirectionality index was then computed as follows:

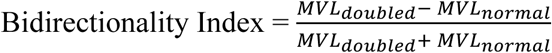

### Cue modulation measures

To determine the extent to which HD/LM-HD cells incorporated the second cue card in the cue duplication session, we fit a bidirectional von Mises function (two peaks or troughs separated by 180°) to each cell’s HD tuning curve from the cue duplication session (LaChance et al., 2022). Trough fits were used only for POR LM-HD cells with maximal firing directions oriented away from the cue card, and MEC/PaS HD cells were fit with upright von Mises functions as they did not exhibit trough tuning. Modulation by the south cue (indicated below by subscript S) was calculated by finding the von Mises peak or trough that was closest to the cell’s session 1 peak or trough, then computing the firing rate difference between that peak (P_S_) or trough (T_S_) and the minimum (Min_fit_) or maximum (Max_fit_) of the fit curve, respectively. This firing rate difference was transformed into a modulation index (MI_S_) by dividing it by the maximum firing rate of the fit curve (Max_fit_):

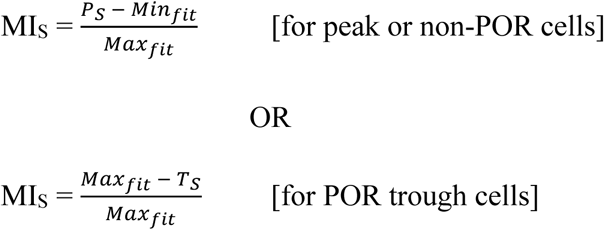

The MI for the north cue was calculated by performing the same computation on the peak or trough 180° opposite.

### Linear mixed model for assessing impact of task experience on cue modulation

To assess whether the impact of visual cue manipulations on HD/LM-HD cells could be explained by the amount of task experience rather than the assigned brain area, we used a linear mixed model (Python package *statsmodels*, function *mixedlm*) to compute the relationship between task experience (defined as number of days an animal had been exposed to the standard task) and either absolute preferred direction shift (for cue rotation sessions) or change in bidirectionality index (for the cue duplication session) while allowing each brain area to have a random slope and intercept.

### Shuffling procedure for cell classifications

Each cell’s spike train was randomly shifted by at least 30 s, with time points beyond the end of the session wrapped to the beginning, to offset the spike data from the behavioral data while maintaining its temporal structure. Relevant tuning scores were then computed based on the shifted spike train. This procedure was repeated 400 times for each cell, and a within-cell 99^th^ percentile cutoff was used to determine tuning significance for individual cells.

### Statistics

Statistical analyses were performed using Python code. All tests were two-sided (except for GLM classifier cross-validation comparisons (Hardcastle et al., 2017; LaChance et al., 2019)) and used an α level of 0.05. Within-cell comparisons across multiple conditions were assessed using a one-way repeated-measures ANOVA. If samples violated sphericity (assessed using Mauchly’s test), we applied a Greenhouse–Geisser correction. Unpaired comparisons were assessed using a one-way ANOVA. *Post hoc* pairwise comparisons were performed using Bonferroni-corrected *t*-tests (Python package Pingouin). Shifts in HD cell preferred firing directions were quantified using a Rayleigh test to confirm clustering of shifts and a circular V-test to test if shifts clustered near a predicted value (Batschelet, 1981). Angle values in the text are given ± circular standard deviation, while non-angle values are given ± standard deviation.

## References

Agster KL, Burwell RD (2013) Hippocampal and subicular efferents and afferents of the perirhinal, postrhinal, and entorhinal cortices of the rat. Behav Brain Res 254:50–64.

Alexander AS, Robinson JC, Stern CE, Hasselmo ME (2023) Gated transformations from egocentric to allocentric reference frames involving retrosplenial cortex, entorhinal cortex, and hippocampus. Hippocampus 33:465–487.

Banino A et al. (2018) Vector-based navigation using grid-like representations in artificial agents. Nature 557:429–433.

Barry C, Lever C, Hayman R, Hartley T, Burton S, O’Keefe J, Jeffery K, Burgess N (2006) The boundary vector cell model of place cell firing and spatial memory. Rev Neurosci 17:71–97.

Batschelet E (1981) Circular Statistics in Biology: Academic Press.

Beltramo R, Scanziani M (2019) A collicular visual cortex: Neocortical space for an ancient midbrain visual structure. Science 363:64–69.

Bicanski A, Burgess N (2018) A neural-level model of spatial memory and imagery. Elife 7.

Boccara CN, Nardin M, Stella F, O’Neill J, Csicsvari J (2019) The entorhinal cognitive map is attracted to goals. Science 363:1443–1447.

Boccara CN, Sargolini F, Thoresen VH, Solstad T, Witter MP, Moser EI, Moser MB (2010) Grid cells in pre- and parasubiculum. Nat Neurosci 13:987–994.

Breese CR, Hampson RE, Deadwyler SA (1989) Hippocampal place cells: stereotypy and plasticity. J Neurosci 9:1097–1111.

Bucci DJ, Burwell RD (2004) Deficits in attentional orienting following damage to the perirhinal or postrhinal cortices. Behav Neurosci 118:1117–1122.

Burgess N, Cacucci F, Lever C, O’keefe J (2005) Characterizing multiple independent behavioral correlates of cell firing in freely moving animals. Hippocampus 15:149–153.

Burwell RD (2000) The parahippocampal region: corticocortical connectivity. Ann N Y Acad Sci 911:25–42.

Burwell RD (2001) Borders and cytoarchitecture of the perirhinal and postrhinal cortices in the rat. J Comp Neurol 437:17–41.

Burwell RD, Amaral DG (1998a) Perirhinal and postrhinal cortices of the rat: interconnectivity and connections with the entorhinal cortex. J Comp Neurol 391:293–321.

Burwell RD, Amaral DG (1998b) Cortical afferents of the perirhinal, postrhinal, and entorhinal cortices of the rat. J Comp Neurol 398:179–205.

Burwell RD, Saddoris MP, Bucci DJ, Wiig KA (2004) Corticohippocampal contributions to spatial and contextual learning. J Neurosci 24:3826–3836.

Bush D, Barry C, Manson D, Burgess N (2015) Using Grid Cells for Navigation. Neuron 87:507–520.

Butler WN, Hardcastle K, Giocomo LM (2019) Remembered reward locations restructure entorhinal spatial maps. Science 363:1447–1452.

Carpenter F, Manson D, Jeffery K, Burgess N, Barry C (2015) Grid cells form a global representation of connected environments. Curr Biol 25:1176–1182.

Clark BJ, LaChance PA, Winter SS, Mehlman ML, Butler W, LaCour A, Taube JS (2024) Comparison of head direction cell firing characteristics across thalamo-parahippocampal circuitry. Hippocampus 34:168–196.

Danielson NB, Zaremba JD, Kaifosh P, Bowler J, Ladow M, Losonczy A (2016) Sublayer-Specific Coding Dynamics during Spatial Navigation and Learning in Hippocampal Area CA1. Neuron 91:652–665.

Davies M, Machin PE, Sanderson DJ, Pearce JM, Aggleton JP (2007) Neurotoxic lesions of the rat perirhinal and postrhinal cortices and their impact on biconditional visual discrimination tasks. Behav Brain Res 176:274–283.

Derdikman D, Whitlock JR, Tsao A, Fyhn M, Hafting T, Moser MB, Moser EI (2009) Fragmentation of grid cell maps in a multicompartment environment. Nat Neurosci 12:1325–1332.

Diehl GW, Hon OJ, Leutgeb S, Leutgeb JK (2017) Grid and Nongrid Cells in Medial Entorhinal Cortex Represent Spatial Location and Environmental Features with Complementary Coding Schemes. Neuron 94:83–92.e86.

Doan TP, Lagartos-Donate MJ, Nilssen ES, Ohara S, Witter MP (2019) Convergent Projections from Perirhinal and Postrhinal Cortices Suggest a Multisensory Nature of Lateral, but Not Medial, Entorhinal Cortex. Cell Rep 29:617–627.e617.

Dupret D, O’Neill J, Pleydell-Bouverie B, Csicsvari J (2010) The reorganization and reactivation of hippocampal maps predict spatial memory performance. Nat Neurosci 13:995–1002.

Duvelle É, Grieves RM, Hok V, Poucet B, Arleo A, Jeffery KJ, Save E (2019) Insensitivity of Place Cells to the Value of Spatial Goals in a Two-Choice Flexible Navigation Task. J Neurosci 39:2522–2541.

Edvardsen V, Bicanski A, Burgess N (2020) Navigating with grid and place cells in cluttered environments. Hippocampus 30:220–232.

Eichenbaum H, Yonelinas AP, Ranganath C (2007) The medial temporal lobe and recognition memory. Annu Rev Neurosci 30:123–152.

Erdem UM, Hasselmo M (2012) A goal-directed spatial navigation model using forward trajectory planning based on grid cells. Eur J Neurosci 35:916–931.

Erdem UM, Hasselmo ME (2014) A biologically inspired hierarchical goal directed navigation model. J Physiol Paris 108:28–37.

Fyhn M, Molden S, Hollup S, Moser MB, Moser E (2002) Hippocampal neurons responding to first-time dislocation of a target object. Neuron 35:555–566.

Gallistel CR (1990) The organization of learning. Cambridge, Mass.: MIT Press.

Gofman X, Tocker G, Weiss S, Boccara CN, Lu L, Moser MB, Moser EI, Morris G, Derdikman D (2019) Dissociation between Postrhinal Cortex and Downstream Parahippocampal Regions in the Representation of Egocentric Boundaries. Curr Biol 29:2751–2757.e2754.

Hafting T, Fyhn M, Molden S, Moser MB, Moser EI (2005) Microstructure of a spatial map in the entorhinal cortex. Nature 436:801–806.

Hales JB, Schlesiger MI, Leutgeb JK, Squire LR, Leutgeb S, Clark RE (2014) Medial entorhinal cortex lesions only partially disrupt hippocampal place cells and hippocampus-dependent place memory. Cell Rep 9:893–901.

Hardcastle K, Maheswaranathan N, Ganguli S, Giocomo LM (2017) A Multiplexed, Heterogeneous, and Adaptive Code for Navigation in Medial Entorhinal Cortex. Neuron 94:375–387.e377.

Hok V, Lenck-Santini PP, Save E, Gaussier P, Banquet JP, Poucet B (2007a) A test of the time estimation hypothesis of place cell goal-related activity. J Integr Neurosci 6:367–378.

Hok V, Lenck-Santini PP, Roux S, Save E, Muller RU, Poucet B (2007b) Goal-related activity in hippocampal place cells. J Neurosci 27:472–482.

Hollup SA, Molden S, Donnett JG, Moser MB, Moser EI (2001) Accumulation of hippocampal place fields at the goal location in an annular watermaze task. J Neurosci 21:1635–1644.

Jeffery KJ, Gilbert A, Burton S, Strudwick A (2003) Preserved performance in a hippocampal-dependent spatial task despite complete place cell remapping. Hippocampus 13:175–189.

Kaufman AM, Geiller T, Losonczy A (2020) A Role for the Locus Coeruleus in Hippocampal CA1 Place Cell Reorganization during Spatial Reward Learning. Neuron 105:1018–1026.e1014.

Klatzky RL (1998) Allocentric and Egocentric Spatial Representations: Definitions, Distinctions, and Interconnections. In: Spatial Cognition (Freksa C, Habel C, Wender KF, eds). Berlin, Heidelberg: Springer.

Knierim JJ, Neunuebel JP, Deshmukh SS (2014) Functional correlates of the lateral and medial entorhinal cortex: objects, path integration and local-global reference frames. Philos Trans R Soc Lond B Biol Sci 369:20130369.

Kobayashi T, Nishijo H, Fukuda M, Bures J, Ono T (1997) Task-dependent representations in rat hippocampal place neurons. J Neurophysiol 78:597–613.

Kobayashi T, Tran AH, Nishijo H, Ono T, Matsumoto G (2003) Contribution of hippocampal place cell activity to learning and formation of goal-directed navigation in rats. Neuroscience 117:1025–1035.

Koganezawa N, Gisetstad R, Husby E, Doan TP, Witter MP (2015) Excitatory Postrhinal Projections to Principal Cells in the Medial Entorhinal Cortex. J Neurosci 35:15860–15874.

Krupic J, Bauza M, Burton S, O’Keefe J (2018) Local transformations of the hippocampal cognitive map. Science 359:1143–1146.

Krupic J, Bauza M, Burton S, Barry C, O’Keefe J (2015) Grid cell symmetry is shaped by environmental geometry. Nature 518:232–235.

Kubie JL, Fenton AA (2012) Linear look-ahead in conjunctive cells: an entorhinal mechanism for vector-based navigation. Front Neural Circuits 6:20.

LaChance PA, Taube JS (2023a) A model for transforming egocentric views into goal-directed behavior. Hippocampus 33:488–504.

LaChance PA, Taube JS (2023b) Geometric determinants of the postrhinal egocentric spatial map. Curr Biol 33:1728–1743.e1727.

LaChance PA, Taube JS (2024) The Anterior Thalamus Preferentially Drives Allocentric But Not Egocentric Orientation Tuning in Postrhinal Cortex. J Neurosci 44.

LaChance PA, Hasselmo ME (2024) Distinct codes for environment structure and symmetry in postrhinal and retrosplenial cortices. Nat Commun 15:8025.

LaChance PA, Todd TP, Taube JS (2019) A sense of space in postrhinal cortex. Science 365.

LaChance PA, Graham J, Shapiro BL, Morris AJ, Taube JS (2022) Landmark-modulated directional coding in postrhinal cortex. Sci Adv 8:eabg8404.

Lee I, Griffin AL, Zilli EA, Eichenbaum H, Hasselmo ME (2006) Gradual translocation of spatial correlates of neuronal firing in the hippocampus toward prospective reward locations. Neuron 51:639–650.

Lever C, Burton S, Jeewajee A, O’Keefe J, Burgess N (2009) Boundary vector cells in the subiculum of the hippocampal formation. J Neurosci 29:9771–9777.

Liu P, Bilkey DK (2002) The effects of NMDA lesions centered on the postrhinal cortex on spatial memory tasks in the rat. Behav Neurosci 116:860–873.

Mamad O, Stumpp L, McNamara HM, Ramakrishnan C, Deisseroth K, Reilly RB, Tsanov M (2017) Place field assembly distribution encodes preferred locations. PLoS Biol 15:e2002365.

Mathis A, Mamidanna P, Cury KM, Abe T, Murthy VN, Mathis MW, Bethge M (2018) DeepLabCut: markerless pose estimation of user-defined body parts with deep learning. Nat Neurosci 21:1281–1289.

McNaughton BL, Battaglia FP, Jensen O, Moser EI, Moser MB (2006) Path integration and the neural basis of the ’cognitive map’. Nat Rev Neurosci 7:663–678.

Morris RG, Garrud P, Rawlins JN, O’Keefe J (1982) Place navigation impaired in rats with hippocampal lesions. Nature 297:681–683.

Moser EI, Kropff E, Moser MB (2008) Place cells, grid cells, and the brain’s spatial representation system. Annu Rev Neurosci 31:69–89.

Naber PA, Witter MP, Lopes da Silva FH (2001) Evidence for a direct projection from the postrhinal cortex to the subiculum in the rat. Hippocampus 11:105–117.

Naber PA, Caballero-Bleda M, Jorritsma-Byham B, Witter MP (1997) Parallel input to the hippocampal memory system through peri- and postrhinal cortices. Neuroreport 8:2617–2621.

Nagahara AH, Otto T, Gallagher M (1995) Entorhinal-perirhinal lesions impair performance of rats on two versions of place learning in the Morris water maze. Behav Neurosci 109:3–9.

Neunuebel JP, Yoganarasimha D, Rao G, Knierim JJ (2013) Conflicts between local and global spatial frameworks dissociate neural representations of the lateral and medial entorhinal cortex. J Neurosci 33:9246–9258.

Nyberg N, Duvelle É, Barry C, Spiers HJ (2022) Spatial goal coding in the hippocampal formation. Neuron 110:394–422.

O’Keefe J (1976) Place units in the hippocampus of the freely moving rat. Exp Neurol 51:78–109.

O’Keefe J, Krupic J (2021) Do hippocampal pyramidal cells respond to nonspatial stimuli? Physiol Rev 101:1427–1456.

Ormond J, O’Keefe J (2022) Hippocampal place cells have goal-oriented vector fields during navigation. Nature 607:741–746.

Pachitariu M, Sridhar S, Stringer C (2023) Solving the spike sorting problem with Kilosort. bioRxiv.

Packard MG, McGaugh JL (1996) Inactivation of hippocampus or caudate nucleus with lidocaine differentially affects expression of place and response learning. Neurobiol Learn Mem 65:65–72.

Peng J-J, Throm B, Jazi MN, Yen T-Y, Monyer H, Allen K (2023) Grid cells perform path integration in multiple reference frames during self-motion-based navigation. bioRxiv:2023.2012.2021.572857.

Pfeiffer BE, Foster DJ (2013) Hippocampal place-cell sequences depict future paths to remembered goals. Nature 497:74–79.

Recce M, Harris KD (1996) Memory for places: a navigational model in support of Marr’s theory of hippocampal function. Hippocampus 6:735–748.

Redish AD, Touretzky DS (1998) The role of the hippocampus in solving the Morris water maze. Neural Comput 10:73–111.

Rossier J, Kaminsky Y, Schenk F, Bures J (2000) The place preference task: a new tool for studying the relation between behavior and place cell activity in rats. Behav Neurosci 114:273–284.

Sanguinetti-Scheck JI, Brecht M (2020) Home, head direction stability, and grid cell distortion. J Neurophysiol 123:1392–1406.

Sarel A, Finkelstein A, Las L, Ulanovsky N (2017) Vectorial representation of spatial goals in the hippocampus of bats. Science 355:176–180.

Sargolini F, Fyhn M, Hafting T, McNaughton BL, Witter MP, Moser MB, Moser EI (2006) Conjunctive representation of position, direction, and velocity in entorhinal cortex. Science 312:758–762.

Savelli F, Yoganarasimha D, Knierim JJ (2008) Influence of boundary removal on the spatial representations of the medial entorhinal cortex. Hippocampus 18:1270–1282.

Schmitzer-Torbert N, Jackson J, Henze D, Harris K, Redish AD (2005) Quantitative measures of cluster quality for use in extracellular recordings. Neuroscience 131:1–11.

Skaggs WE, McNaughton BL, Gothard KM, Markus EJ (1997) An Information-Theoretic Approach to Deciphering the Hippocampal Code. In: Adv Neural Inf Process Syst.

Solstad T, Boccara CN, Kropff E, Moser MB, Moser EI (2008) Representation of geometric borders in the entorhinal cortex. Science 322:1865–1868.

Sosa M, Giocomo LM (2021) Navigating for reward. Nat Rev Neurosci 22:472–487.

Spiers HJ, Olafsdottir HF, Lever C (2018) Hippocampal CA1 activity correlated with the distance to the goal and navigation performance. Hippocampus 28:644–658.

Steward O, Scoville SA (1976) Cells of origin of entorhinal cortical afferents to the hippocampus and fascia dentata of the rat. J Comp Neurol 169:347–370.

Taube JS (1995) Head direction cells recorded in the anterior thalamic nuclei of freely moving rats. J Neurosci 15:70–86.

Taube JS, Muller RU, Ranck JB (1990) Head-direction cells recorded from the postsubiculum in freely moving rats. I. Description and quantitative analysis. J Neurosci 10:420–435.

Tomás Pereira I, Agster KL, Burwell RD (2016) Subcortical connections of the perirhinal, postrhinal, and entorhinal cortices of the rat. I. afferents. Hippocampus 26:1189–1212.

Turi GF, Li WK, Chavlis S, Pandi I, O’Hare J, Priestley JB, Grosmark AD, Liao Z, Ladow M, Zhang JF, Zemelman BV, Poirazi P, Losonczy A (2019) Vasoactive Intestinal Polypeptide-Expressing Interneurons in the Hippocampus Support Goal-Oriented Spatial Learning. Neuron 101:1150–1165.e1158.

van Groen T, Wyss JM (1990) The connections of presubiculum and parasubiculum in the rat. Brain Res 518:227–243.

Vollan AZ, Gardner RJ, Moser MB, Moser EI (2025) Left-right-alternating theta sweeps in entorhinal-hippocampal maps of space. Nature 639:995–1005.

Vöröslakos M, Miyawaki H, Royer S, Diba K, Yoon E, Petersen PC, Buzsáki G (2021) 3D-printed Recoverable Microdrive and Base Plate System for Rodent Electrophysiology. Bio Protoc 11:e4137.

Wang C, Chen X, Lee H, Deshmukh SS, Yoganarasimha D, Savelli F, Knierim JJ (2018) Egocentric coding of external items in the lateral entorhinal cortex. Science 362:945–949.

Winter SS, Clark BJ, Taube JS (2015) Spatial navigation. Disruption of the head direction cell network impairs the parahippocampal grid cell signal. Science 347:870–874.

Witter MP, Naber PA, van Haeften T, Machielsen WC, Rombouts SA, Barkhof F, Scheltens P, Lopes da Silva FH (2000) Cortico-hippocampal communication by way of parallel parahippocampal-subicular pathways. Hippocampus 10:398–410.

Xiao Z, Lin K, Fellous JM (2020) Conjunctive reward-place coding properties of dorsal distal CA1 hippocampus cells. Biol Cybern 114:285–301.

Xu H, Baracskay P, O’Neill J, Csicsvari J (2019) Assembly Responses of Hippocampal CA1 Place Cells Predict Learned Behavior in Goal-Directed Spatial Tasks on the Radial Eight-Arm Maze. Neuron 101:119–132.e114.

Zaremba JD, Diamantopoulou A, Danielson NB, Grosmark AD, Kaifosh PW, Bowler JC, Liao Z, Sparks FT, Gogos JA, Losonczy A (2017) Impaired hippocampal place cell dynamics in a mouse model of the 22q11.2 deletion. Nat Neurosci 20:1612–1623.

Zhang GR, Cao H, Kong L, O’Brien J, Baughns A, Jan M, Zhao H, Wang X, Lu XG, Cook RG, Geller AI (2010) Identified circuit in rat postrhinal cortex encodes essential information for performing specific visual shape discriminations. Proc Natl Acad Sci U S A 107:14478–14483.

