## Supplemental Figures 1-14 for "Cortical dissociation of spatial reference frames during place navigation"

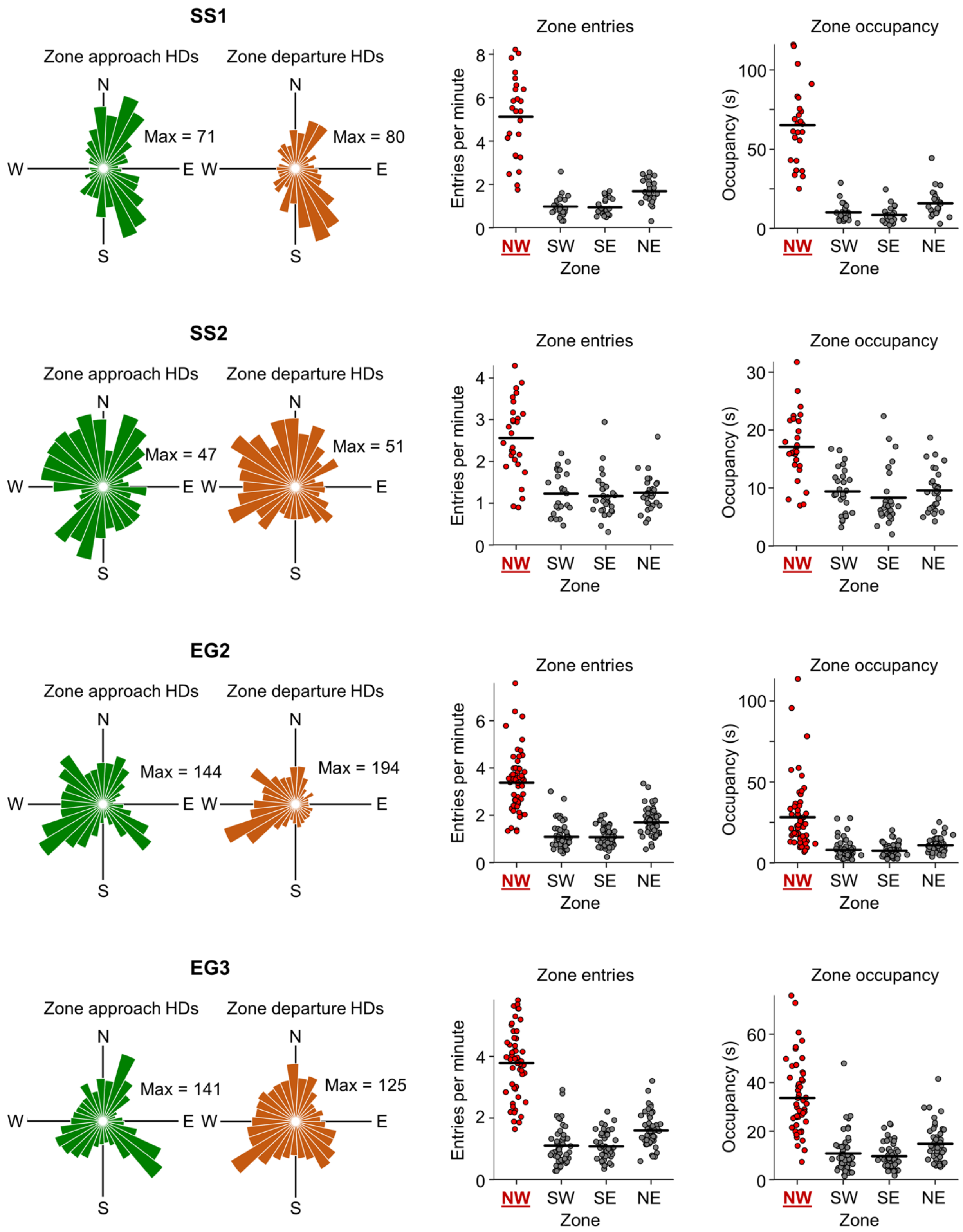


**Supplemental Figure 1. Task performance for individual animals.** *Left column,* polar histograms showing the distribution of head directions for all goal zone approaches (*left*) and departures (*right*) for all standard task sessions performed by each animal. *Middle column*, strip plot showing the number of entries per minute into the goal zone vs. rotationally equivalent zones. *Right column*, same as middle column but showing occupancy time in the goal zone vs. rotationally equivalent zones. Note that all animals tended to enter the goal zone and spend more time there than any of the equivalent zones.

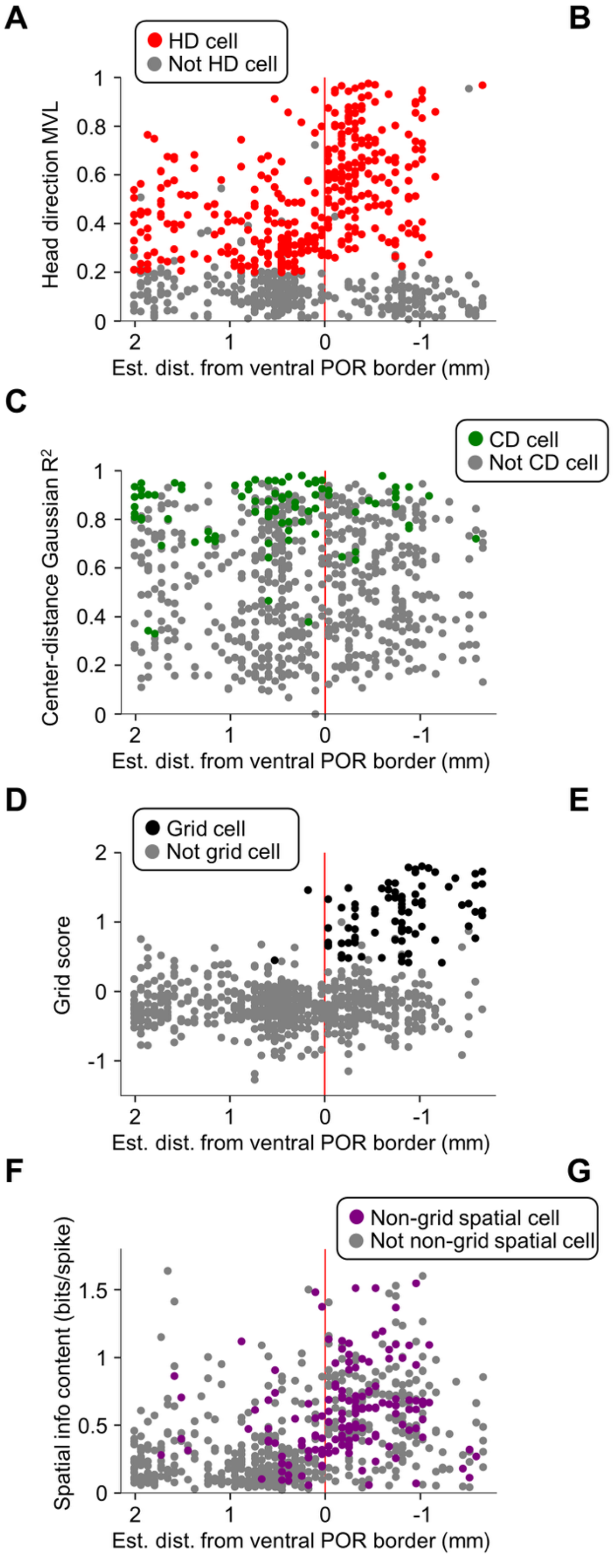


**Supplemental Figure 2. Cell tuning properties relative to estimated anatomical placement.** **A)** Scatter plot of HD tuning curve MVLs compared to estimated distance from ventral POR border. Note the abrupt increase in HD cell MVLs upon crossing the border. **B)** Same as **(A)** but for center-bearing (CB) tuning curve MVLs. Note the decrease in MVLs upon crossing the ventral POR border. **C)** Same as **(A)** but for Gaussian (*left*) or linear (*right*) $R^2$ fit values for center-distance (CD) tuning curves relative to the center of the environment. Note that tuning to the center is roughly equivalent to tuning to the environmental boundaries. **D)** Same as **(A)** but for grid scores. Note the sudden appearance of grid cells upon crossing the border. **E)** Same as **(A)** but for border scores. **F)** Same as **(A)** but for spatial information content. Note the increased spatial information content in MEC/PaS compared to POR. **G)** Same as **(A)** but for theta index based on spike autocorrelograms. Note the abrupt appearance of theta-modulated cells upon crossing the ventral POR border.

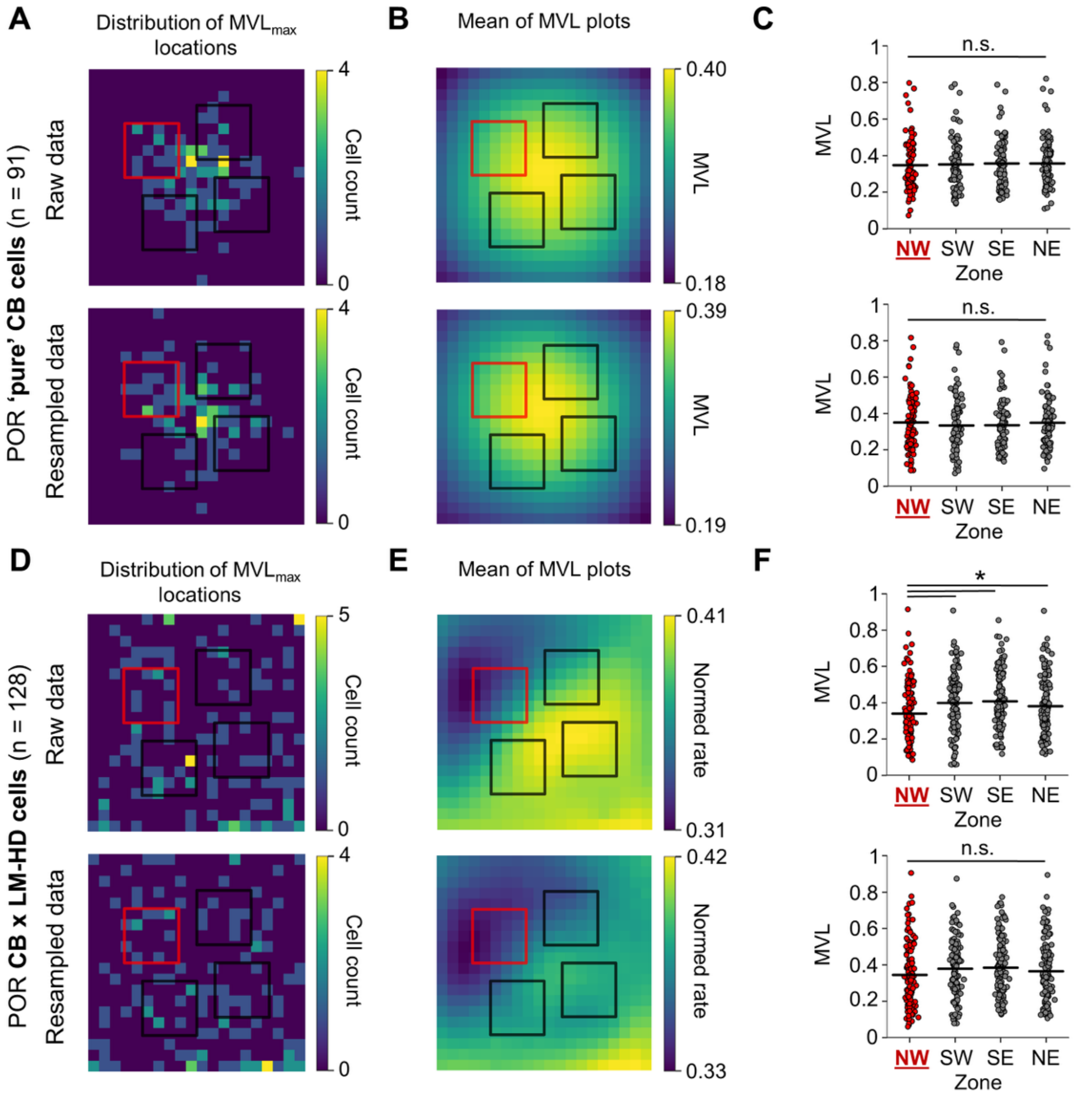


**Supplemental Figure 3. Equivalent zone MVL analyses for POR cells. A)** 2D histogram showing the distribution of $MVL_{max}$ locations for all POR 'pure' CB cells recorded in the baseline session for raw data (*top*) and data that was resampled to match spatial occupancy across the task environment (*bottom*). Overlaid boxes indicate expanded (30 x 30 cm) zones centered on the goal zone (red) and rotationally equivalent zones (black). $MVL_{max}$ locations were not more common in the expanded goal zone compared to the other expanded zones for either raw or resampled data (raw data: $\chi^2(3) = 0.6$, $P = 0.90$; resampled data: $\chi^2(3) = 3.8$, $P = 0.28$). **B)** Mean of MVL plots for all 'pure' CB cells based on raw (*top*) and resampled (*bottom*) data. Note the concentration of high MVLs in the center of the environment. **C)** Strip plots comparing the mean MVL values in the expanded goal zone with the expanded equivalent zones across all 'pure' CB cells for raw (*top*) and resampled (*bottom*) data. No significant differences were observed (raw data: repeated measures ANOVA, $F(3, 270) = 0.82$, $P = 0.48$; resampled data: repeated measures ANOVA, $F(3, 270) = 2.43$, $P = 0.065$). **D)** Same as **(A)** but for all POR CB x LM-HD cells recorded in the baseline session. $MVL_{max}$ frequencies did not differ between the expanded zones (raw data: $\chi^2(3) = 6.26$, $P = 0.10$; resampled data: $\chi^2(3) = 1.0$, $P = 0.80$). **E-F)** Same as **(B-C)** but for CB x LM-HD cells. Note that MVLs were significantly lower on average in the expanded goal zone compared to the expanded equivalent zones based on the raw data, but no differences were found when the data was resampled to remove spatial bias toward the goal zone (raw data: repeated measures ANOVA, $F(3, 381) = 9.26$, $P = 7.83e\text{-}5$; pairwise *t*-tests, NW vs. SW, $t(127) = -4.23$, $P = 1.34e\text{-}4$; NW vs. SE, $t(127) = -4.05$, $P = 2.67e\text{-}4$; NW vs. NE, $t(127) = -3.07$, $P = 7.82e\text{-}3$; resampled data: repeated measures ANOVA, $F(3, 381) = 2.63$, $P = 0.068$).

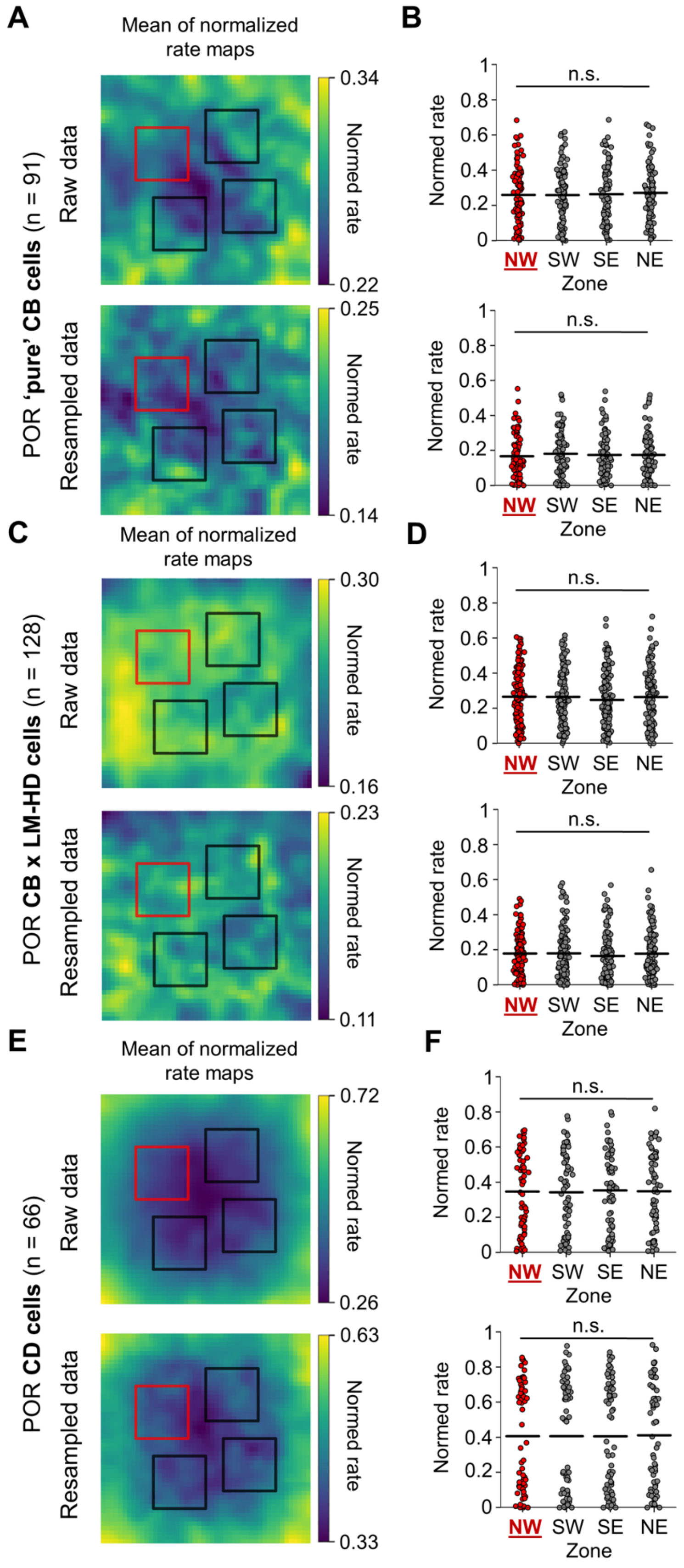
A
Mean of normalized rate maps
POR 'pure' CB cells (n = 91)
Raw data
Resampled data
Normed rate
0.34
0.22
0.25
0.14
B
Normed rate
n.s.
NW SW SE NE
Zone
C
Mean of normalized rate maps
POR CB x LM-HD cells (n = 128)
Raw data
Resampled data
Normed rate
0.30
0.16
0.23
0.11
D
E
Mean of normalized rate maps
POR CD cells (n = 66)
Raw data
Resampled data
Normed rate
0.72
0.26
0.63
0.33
F

**Supplemental Figure 4. Equivalent zone firing rate analyses for POR cells. A)** Mean 2D allocentric firing rate map for all POR 'pure' CB cells recorded in the baseline session based on raw data (*top*) and data that was resampled to match spatial occupancy across the task environment (*bottom*). Overlaid boxes indicate expanded (30 x 30 cm) zones centered on the goal zone (red) and rotationally equivalent zones (black). **B)** Strip plots comparing the mean firing rates in the expanded goal zone with the expanded equivalent zones across all 'pure' CB cells for both raw (*top*) and resampled (*bottom*) data. There were no significant differences between the zones (raw data: repeated measures ANOVA, $F(3, 270) = 0.65$, $P = 0.55$; resampled data: repeated measures ANOVA, $F(3, 270) = 0.74$, $P = 0.51$). **C-D)** Same as **(A-B)** but for POR CB x LM-HD cells. There were no significant firing rate differences between the zones (raw data: repeated measures ANOVA, $F(3, 381) = 1.17$, $P = 0.32$; resampled data: repeated measures ANOVA, $F(3, 381) = 1.05$, $P = 0.37$). **E-F)** Same as **(A-B)** but for POR cells tuned to center-distance (CD) in the baseline session. There were no significant firing rate differences between the zones (raw data: repeated measures ANOVA, $F(3, 195) = 0.23$, $P = 0.84$; resampled data: repeated measures ANOVA, $F(3, 195) = 0.11$, $P = 0.95$).

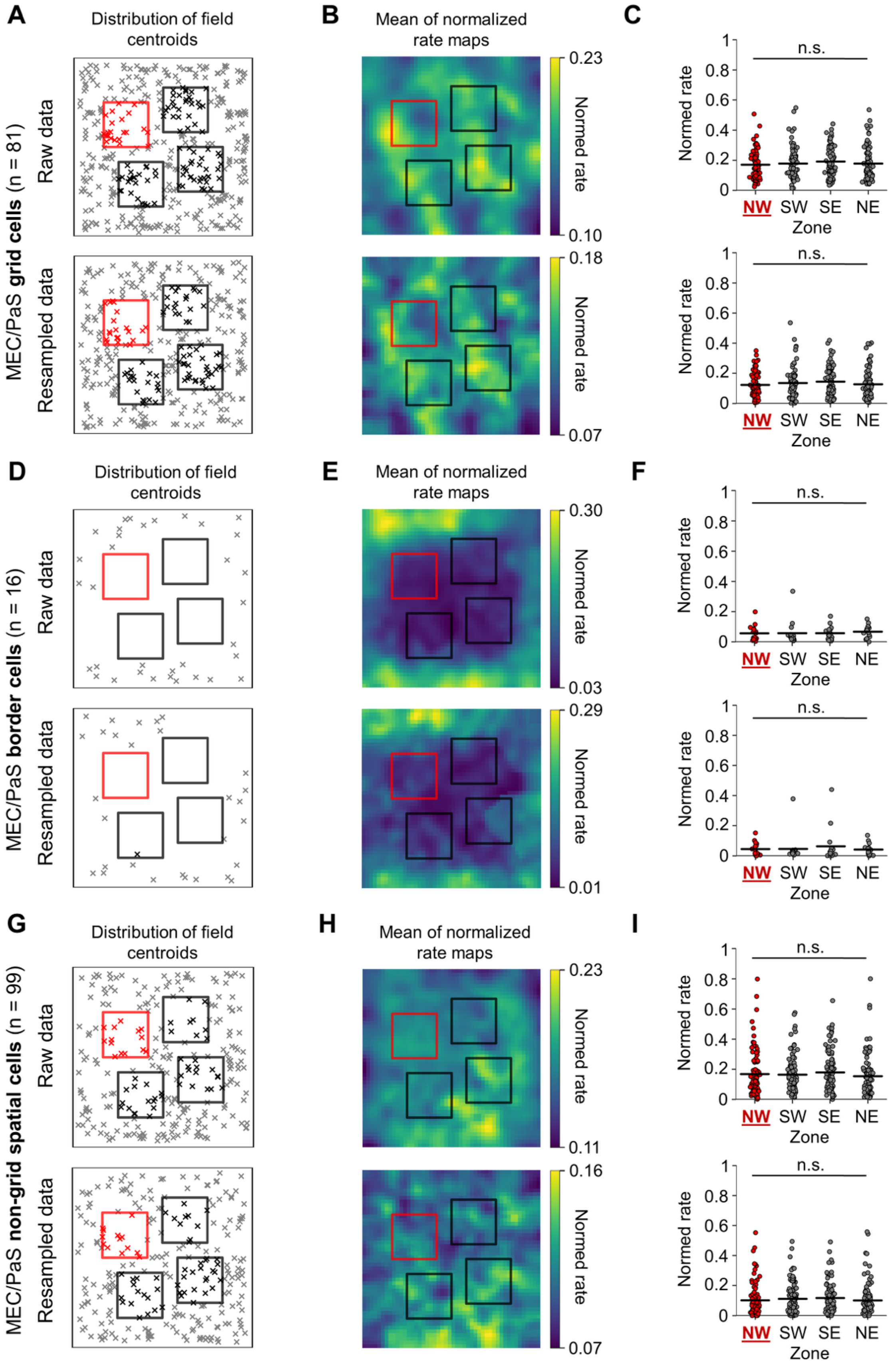

A
Distribution of field centroids
B
Mean of normalized rate maps
C
MEC/PaS grid cells (n = 81)
Raw data
Resampled data
0.23
0.10
0.18
0.07
Normed rate
n.s.
Normed rate
NW SW SE NE
Zone
D
Distribution of field centroids
E
Mean of normalized rate maps
F
MEC/PaS border cells (n = 16)
0.30
0.03
0.29
0.01
G
Distribution of field centroids
H
Mean of normalized rate maps
I
MEC/PaS non-grid spatial cells (n = 99)
0.23
0.11
0.16
0.07

**Supplemental Figure 5. Equivalent zone firing rate analyses for MEC/PaS cells. A)** Scatter plot showing the distribution of firing field centroids for all MEC/PaS grid cells recorded in the baseline session based on raw data (*top*) and data that was resampled to match spatial occupancy across the task environment (*bottom*). Overlaid boxes indicate expanded (30 x 30 cm) zones centered on the goal zone (red) and rotationally equivalent zones (black). Firing field centroids were not more common in the expanded goal zone compared to the expanded equivalent zone based on either raw or resampled data (raw data: $\chi^2(3) = 2.15$, $P = 0.54$; resampled data: $\chi^2(3) = 4.31$, $P = 0.23$). **B)** Mean 2D allocentric firing rate map for all MEC/PaS grid cells recorded in the baseline session based on raw (*top*) and resampled (*bottom*) data. **C)** Strip plots comparing the mean firing rates in the expanded goal zone with the expanded equivalent zones across all grid cells for both raw (*top*) and resampled (*bottom*) data. There were no significant differences between the zones (raw data: repeated measures ANOVA, $F(3, 240) = 0.79$, $P = 0.50$; resampled data: repeated measures ANOVA, $F(3, 240) = 1.36$, $P = 0.26$). **D-F)** Same as **(A-C)** but for MEC/PaS border cells. Border cell firing field centroids did not tend to fall into any of the analyzed zones due to their restricted firing along the walls of the environment. There were also no overall firing rate differences between any of the zones (raw data: repeated measures ANOVA, $F(3, 45) = 0.30$, $P = 0.83$; resampled data: repeated measures ANOVA, $F(3, 45) = 0.31$, $P = 0.82$). **G-I)** Same as **(A-C)** but for MEC/PaS non-grid spatial cells. Firing field centroids were not more common in the expanded goal zone compared to the expanded equivalent zones (raw data: $\chi^2(3) = 2.15$, $P = 0.54$; resampled data: $\chi^2(3) = 5.16$, $P = 0.16$). There were no significant firing rate differences between the zones (raw data: repeated measures ANOVA, $F(3, 294) = 1.39$, $P = 0.25$; resampled data: repeated measures ANOVA, $F(3, 294) = 1.24$, $P = 0.30$).

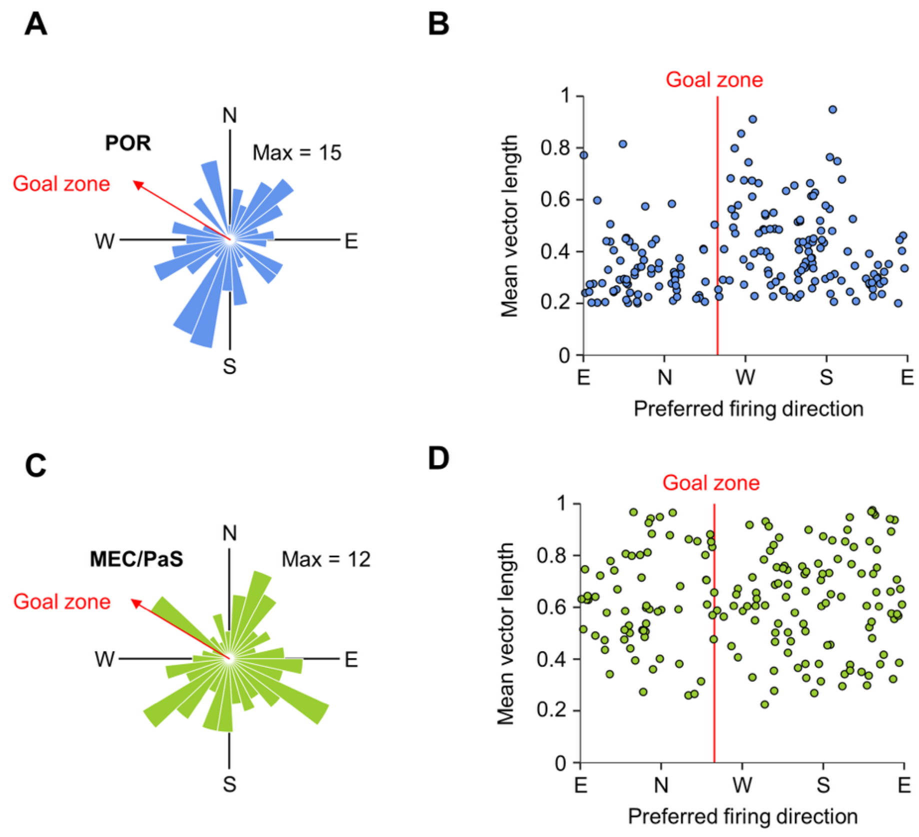


**Supplemental Figure 6. HD cell distributions relative to goal zone. A)** Polar histogram showing the distribution of preferred firing directions for all POR LM-HD cells recorded in the baseline session (N = 192). The red line indicates the general direction of the goal zone (the angle of the goal zone from the environment center). Note that the preferred directions did not cluster around the goal direction. **B)** Scatter plot showing the relationship between preferred firing direction and HD mean vector length for POR LM-HD cells. Note that POR LM-HD cells tend to have higher mean vector lengths in the general direction of the cue card (between 180° and 270°) but not in the direction of the goal zone. **C-D)** Same as **(A-B)** but for MEC/PaS HD cells. Note the overall lack of bias toward the general direction of the goal zone.

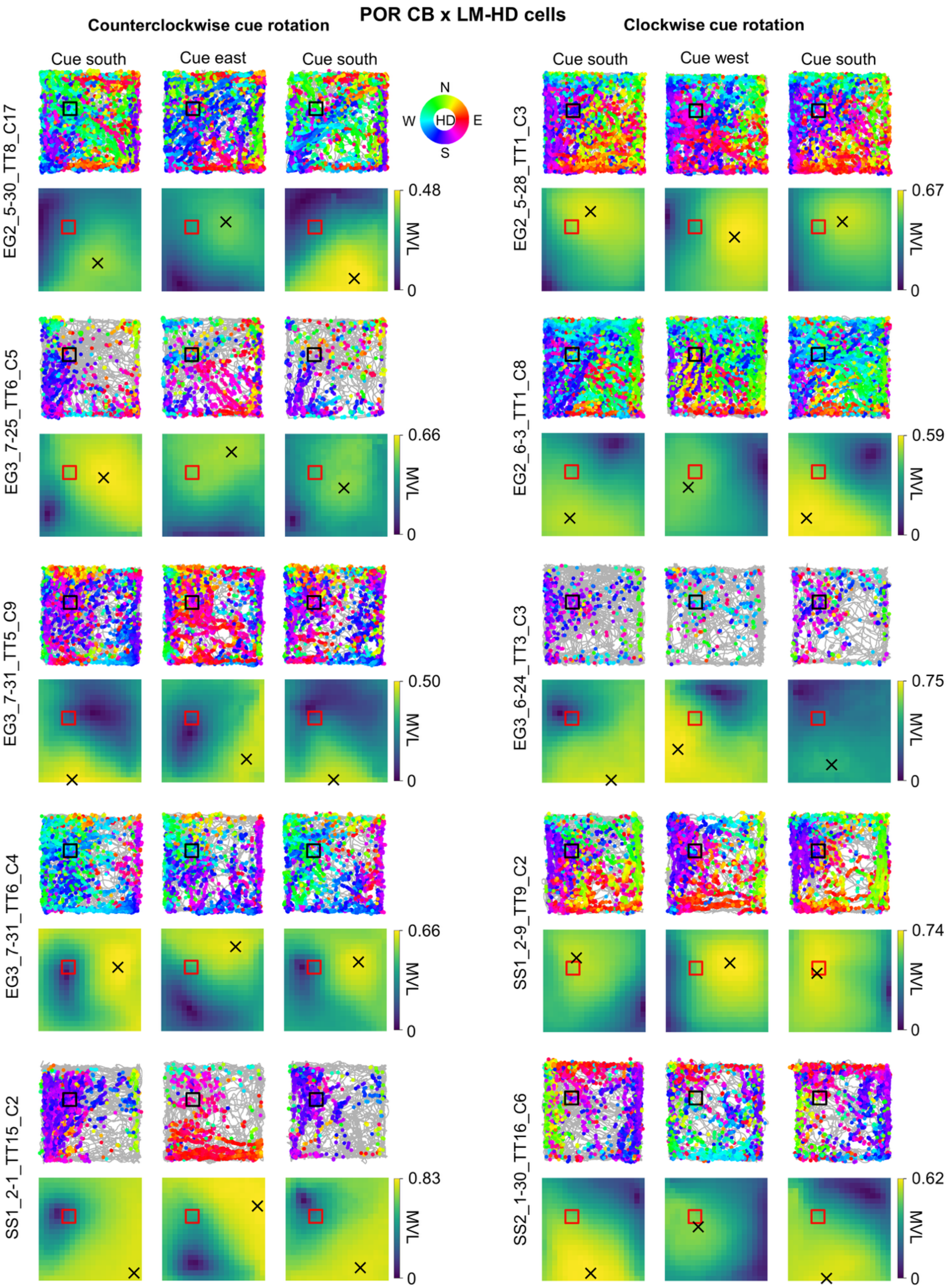

POR CB x LM-HD cells
Counterclockwise cue rotation
Clockwise cue rotation
Cue south
Cue east
Cue south
Cue south
Cue west
Cue south
N
W
HD
E
S
EG2_5-30_TT8_C17
EG3_7-25_TT6_C5
EG3_7-31_TT5_C9
EG3_7-31_TT6_C4
SS1_2-1_TT15_C2
EG2_5-28_TT1_C3
EG2_6-3_TT1_C8
EG3_6-24_TT3_C3
SS1_2-9_TT9_C2
SS2_1-30_TT16_C6
MVL
0.48
0.66
0.50
0.66
0.83
0.67
0.59
0.75
0.74
0.62
0

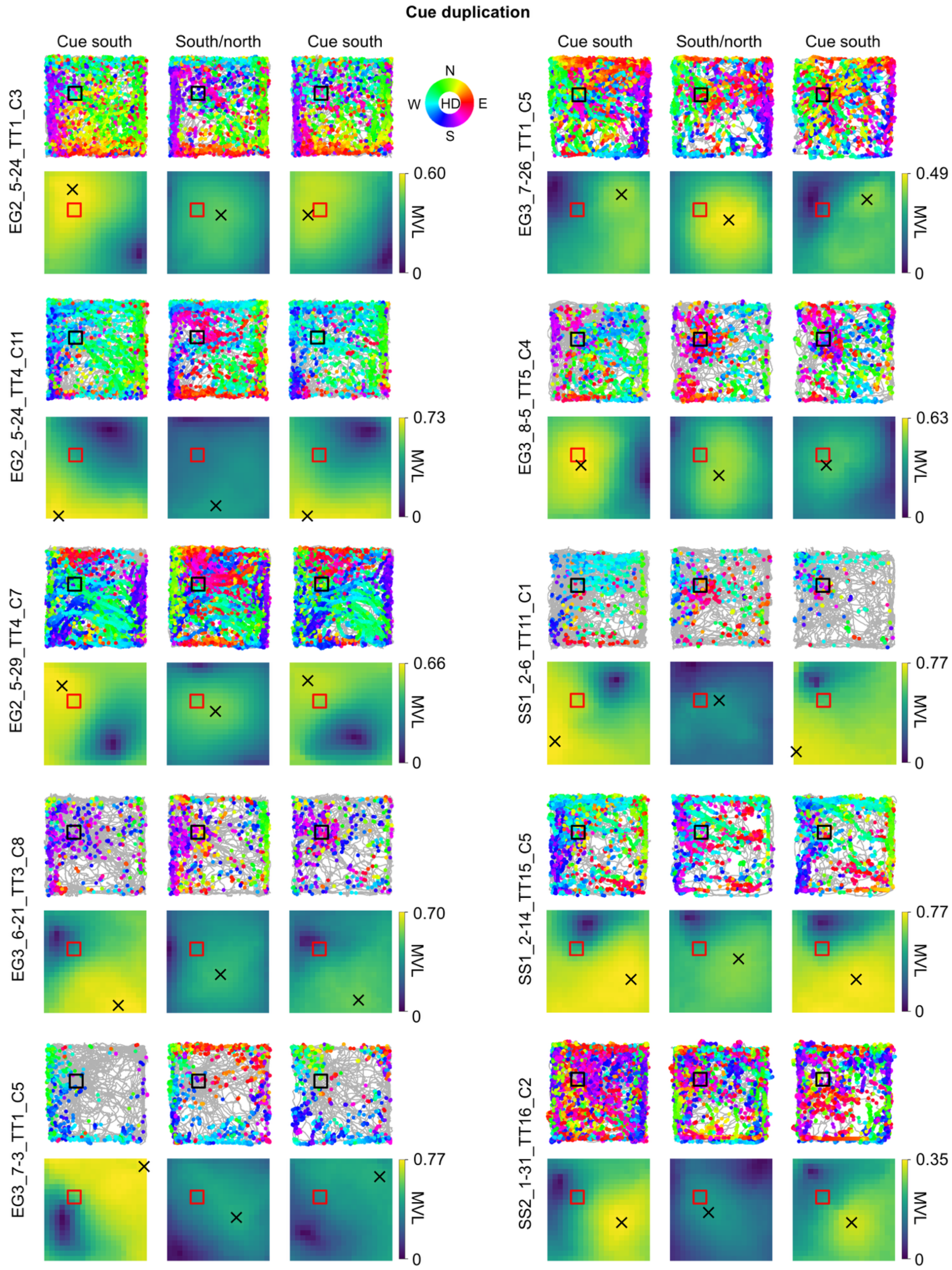
Cue duplication
Cue south
South/north
Cue south
Cue south
South/north
Cue south
N
W
HD
E
S
EG2_5-24_TT1_C3
0.60
MVL
0
EG3_7-26_TT1_C5
0.49
MVL
0
EG2_5-24_TT4_C11
0.73
MVL
0
EG3_8-5_TT5_C4
0.63
MVL
0
EG2_5-29_TT4_C7
0.66
MVL
0
SS1_2-6_TT11_C1
0.77
MVL
0
EG3_6-21_TT3_C8
0.70
MVL
0
SS1_2-14_TT15_C5
0.77
MVL
0
EG3_7-3_TT1_C5
0.77
MVL
0
SS2_1-31_TT16_C2
0.35
MVL
0

**Supplemental Figure 7. Additional POR CB x LM-HD cell examples.** HD-colored path and spike plots (*top row*) and MVL plots (*bottom row*) for five additional POR CB x LM-HD cells recorded in the counterclockwise rotation session, five recorded in the clockwise rotation session, and ten recorded in the cue duplication session.

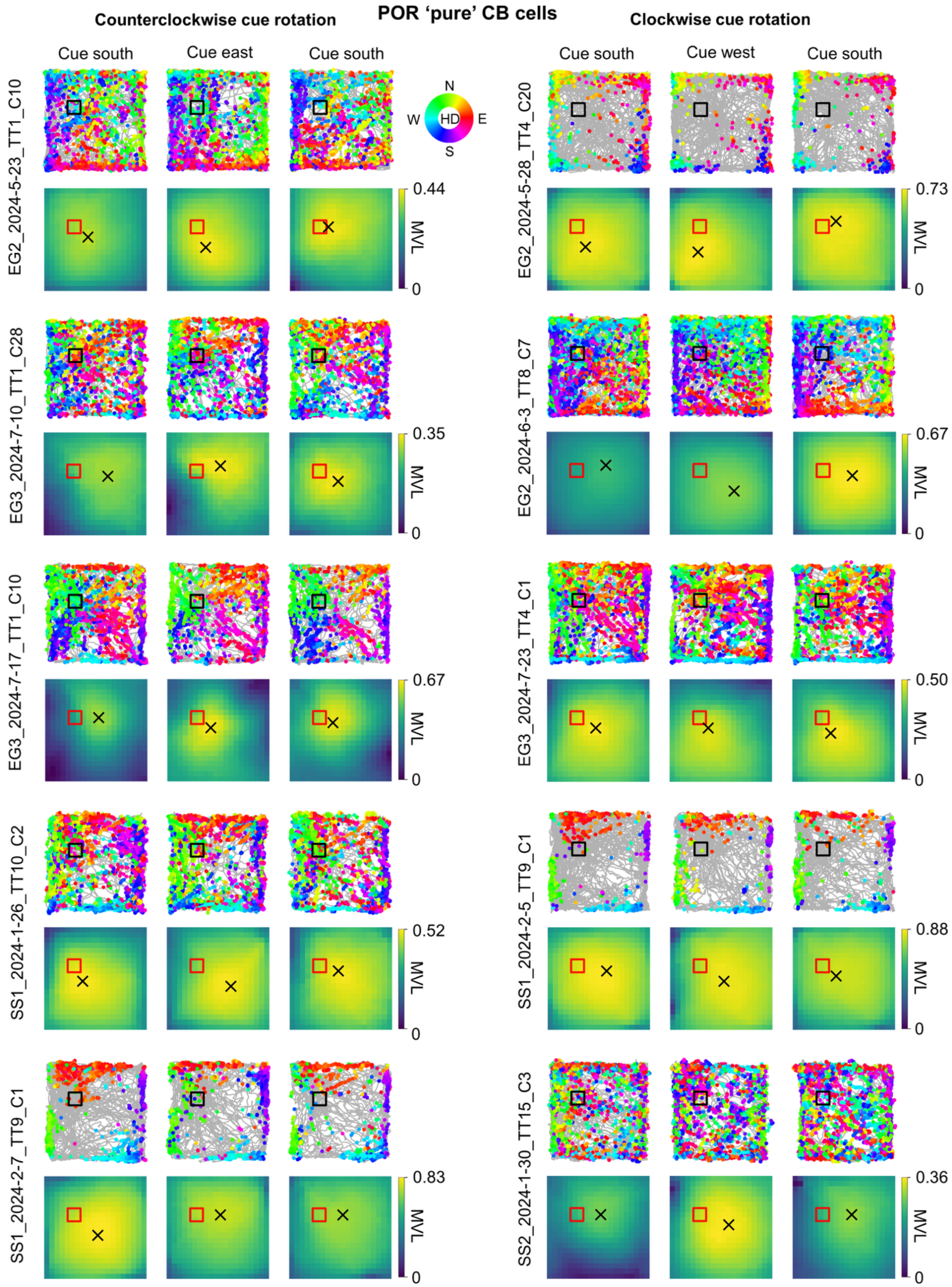
POR ‘pure’ CB cells
Counterclockwise cue rotation
Clockwise cue rotation
Cue south
Cue east
Cue south
Cue south
Cue west
Cue south
N
W
HD
E
S
EG2_2024-5-23_TT1_C10
EG3_2024-7-10_TT1_C28
EG3_2024-7-17_TT1_C10
SS1_2024-1-26_TT10_C2
SS1_2024-2-7_TT9_C1
EG2_2024-5-28_TT4_C20
EG2_2024-6-3_TT8_C7
EG3_2024-7-23_TT4_C1
SS1_2024-2-5_TT9_C1
SS2_2024-1-30_TT15_C3
0.44
0.35
0.67
0.52
0.83
0.73
0.67
0.50
0.88
0.36
MVL
0

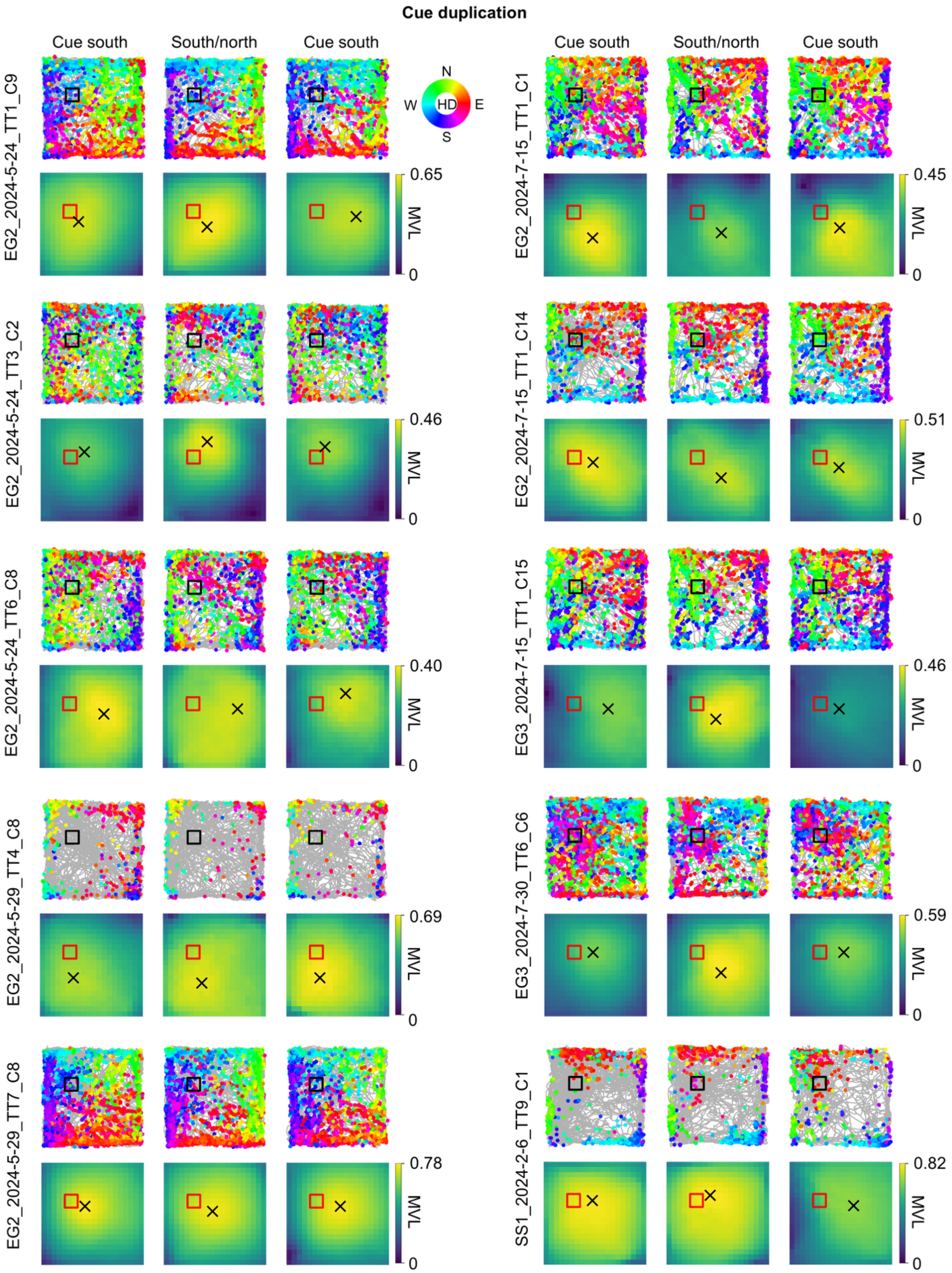

Cue duplication
Cue south
South/north
Cue south
Cue south
South/north
Cue south
N
W
HD
E
S
EG2_2024-5-24_TT1_C9
EG2_2024-7-15_TT1_C1
EG2_2024-5-24_TT3_C2
EG2_2024-7-15_TT1_C14
EG2_2024-5-24_TT6_C8
EG3_2024-7-15_TT1_C15
EG2_2024-5-29_TT4_C8
EG3_2024-7-30_TT6_C6
EG2_2024-5-29_TT7_C8
SS1_2024-2-6_TT9_C1
MVL
0.65
0.45
0.46
0.51
0.40
0.46
0.69
0.59
0.78
0.82
0

**Supplemental Figure 8. Additional POR 'pure' CB cell examples.** HD-colored path and spike plots (*top row*) and MVL plots (*bottom row*) for five additional POR 'pure' CB cells recorded in the counterclockwise rotation session, five recorded in the clockwise rotation session, and ten recorded in the cue duplication session.

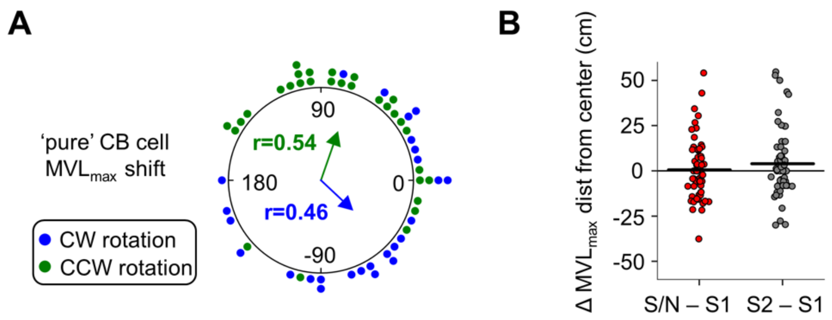


**Supplemental Figure 9. Population response of POR 'pure' CB cells to cue manipulations. A)** Polar dot plot showing the rotation of the $MVL_{max}$ location around the center of the environment for all POR 'pure' CB cells between the initial standard session and the following cue rotation session. Note that the $MVL_{max}$ rotations did not statistically differ from the 90° cue rotation (counterclockwise (n = 30): Rayleigh test, $r = 0.54$, $P = 9.98e-4$; V-test for concentration near 90°, $u = 3.54$, $P = 1.99e-4$; clockwise (n = 26): Rayleigh test, $r = 0.46$, $P = 0.0029$; V-test for concentration near -90°, $u = 2.62$, $P = 0.0044$). **B)** Strip plot showing the change in distance of $MVL_{max}$ locations from the center of the environment for all POR 'pure' CB cells (n = 59 cells) between the initial standard session (S1) and both the cue duplication session (S/N) and the final standard session (S2). Note that, unlike for CB x LM-HD cells, the $MVL_{max}$ locations of 'pure' CB cells did not shift significantly toward the center of the environment in the cue duplication session compared to both standard sessions, likely due to their $MVL_{max}$ locations already being extremely close to the environment center (repeated measures ANOVA, $F(2, 116) = 1.96$, $P = 0.15$).

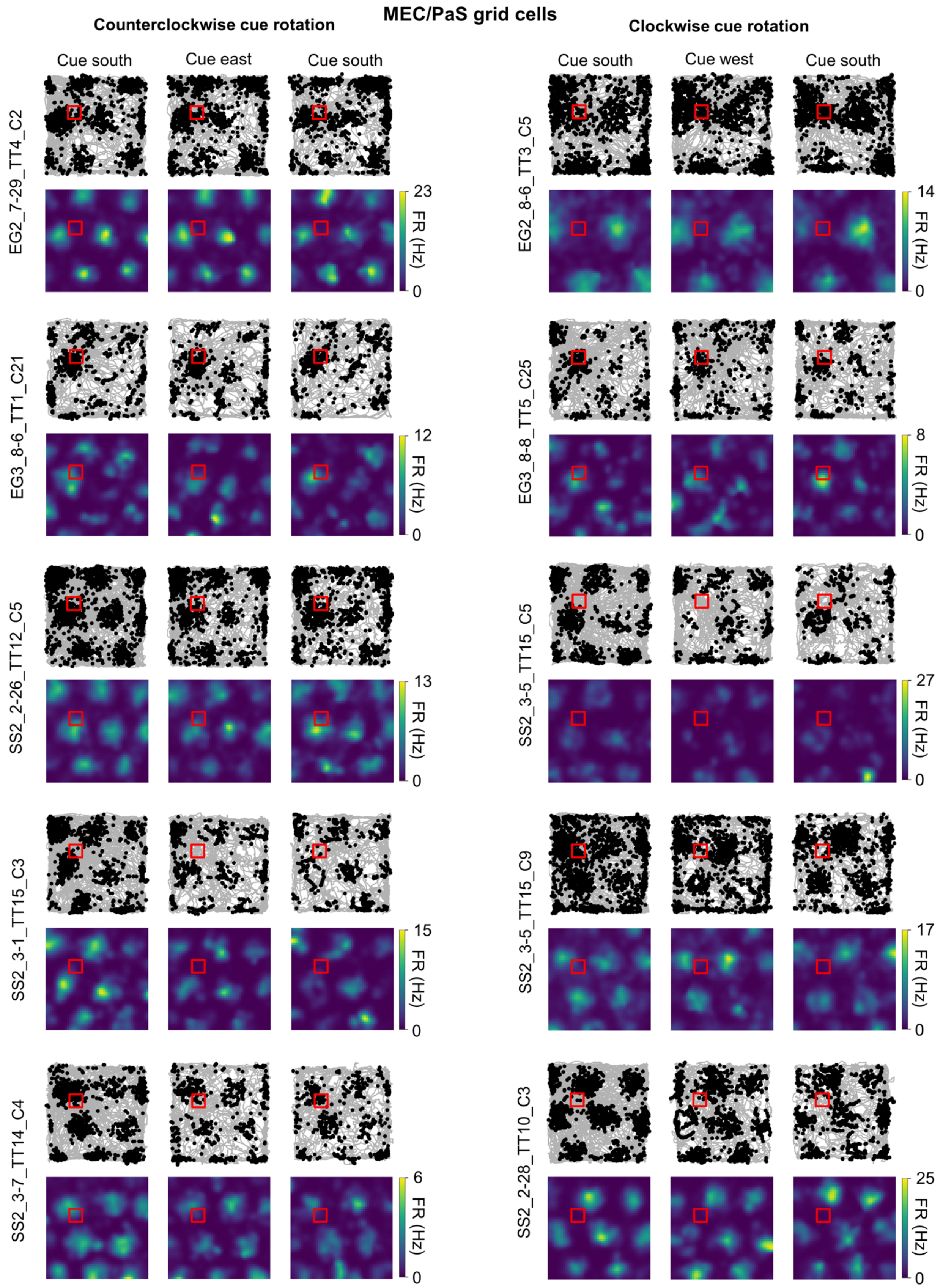
MEC/PaS grid cells
Counterclockwise cue rotation
Clockwise cue rotation
Cue south
Cue east
Cue south
Cue south
Cue west
Cue south
EG2_7-29_TT4_C2
EG3_8-6_TT1_C21
SS2_2-26_TT12_C5
SS2_3-1_TT15_C3
SS2_3-7_TT14_C4
EG2_8-6_TT3_C5
EG3_8-8_TT5_C25
SS2_3-5_TT15_C5
SS2_3-5_TT15_C9
SS2_2-28_TT10_C3
FR (Hz)
23
12
13
15
6
14
8
27
17
25
0

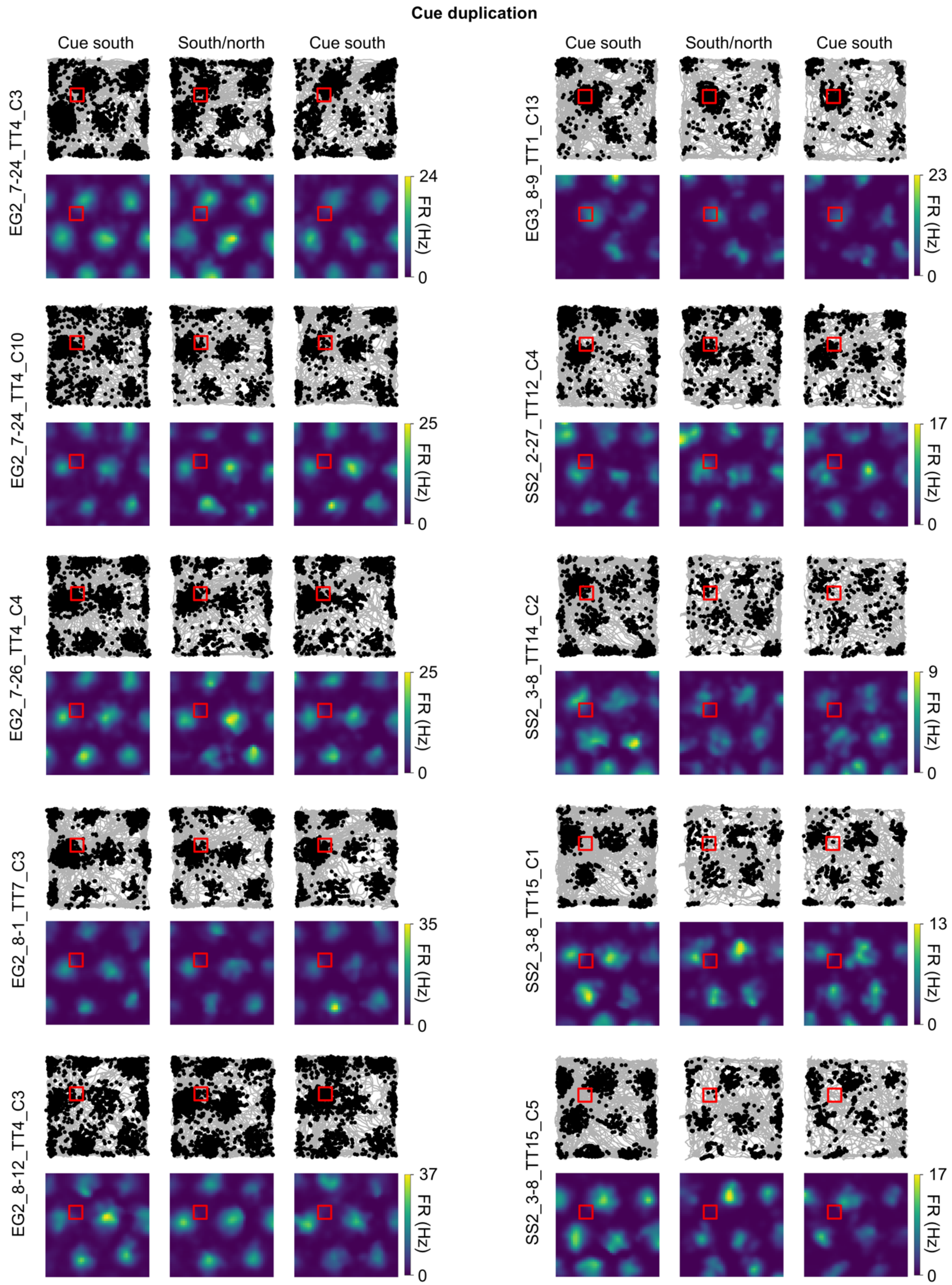

Cue duplication
Cue south
South/north
Cue south
Cue south
South/north
Cue south
EG2_7-24_TT4_C3
24
FR (Hz)
0
EG3_8-9_TT1_C13
23
FR (Hz)
0
EG2_7-24_TT4_C10
25
FR (Hz)
0
SS2_2-27_TT12_C4
17
FR (Hz)
0
EG2_7-26_TT4_C4
25
FR (Hz)
0
SS2_3-8_TT14_C2
9
FR (Hz)
0
EG2_8-1_TT7_C3
35
FR (Hz)
0
SS2_3-8_TT15_C1
13
FR (Hz)
0
EG2_8-12_TT4_C3
37
FR (Hz)
0
SS2_3-8_TT15_C5
17
FR (Hz)
0

**Supplemental Figure 10. Additional MEC/PaS grid cell examples.** Path and spike plots (*top row*) and 2D allocentric firing rate maps (*bottom row*) for five additional MEC/PaS grid cells recorded in the counterclockwise rotation session, five recorded in the clockwise rotation session, and ten recorded in the cue duplication session. Note that grid cells representations were unchanged by the cue manipulations.

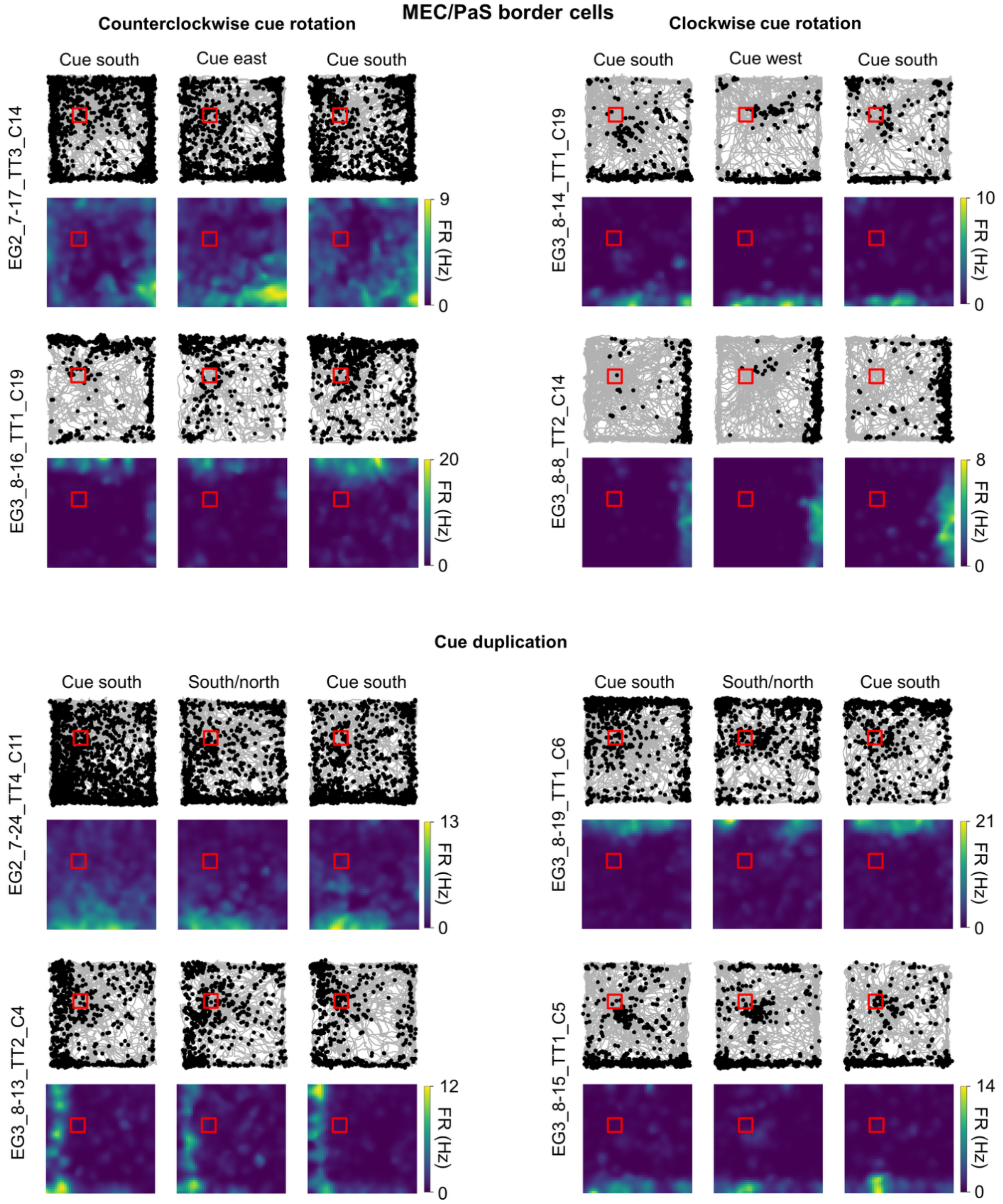


**Supplemental Figure 11. MEC/PaS border cell examples.** Path and spike plots (*top row*) and 2D allocentric firing rate maps (*bottom row*) for two MEC/PaS border cells recorded in the counterclockwise rotation session, two recorded in the clockwise rotation session, and four recorded in the cue duplication session. Note that border cell representations were unchanged by the cue manipulations.

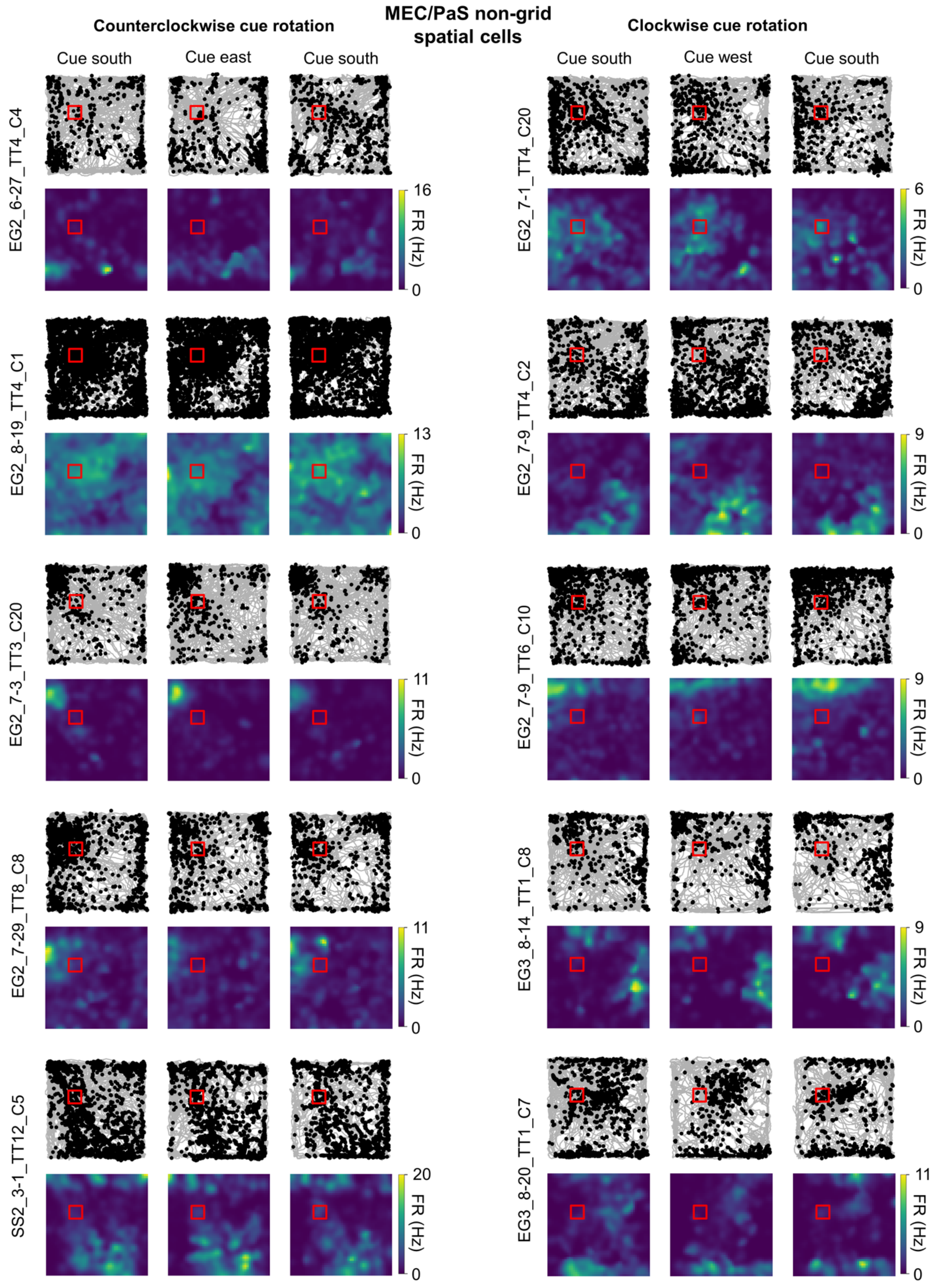
MEC/PaS non-grid spatial cells
Counterclockwise cue rotation
Clockwise cue rotation
Cue south
Cue east
Cue south
Cue south
Cue west
Cue south
EG2_6-27_TT4_C4
EG2_7-1_TT4_C20
EG2_8-19_TT4_C1
EG2_7-9_TT4_C2
EG2_7-3_TT3_C20
EG2_7-9_TT6_C10
EG2_7-29_TT8_C8
EG3_8-14_TT1_C8
SS2_3-1_TT12_C5
EG3_8-20_TT1_C7
FR (Hz)
16
0
6
13
9
11
9
11
9
20
11

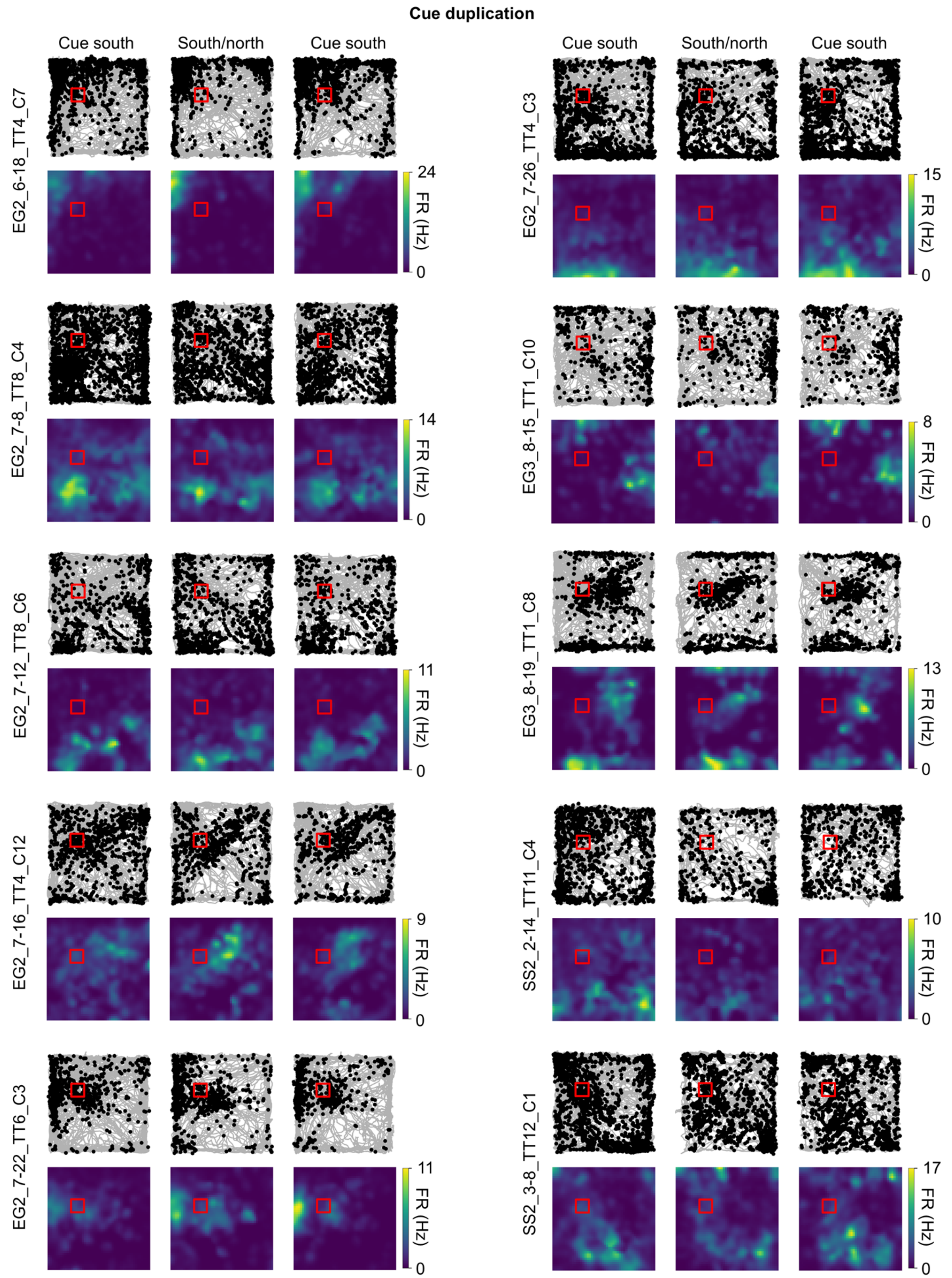
Cue duplication
Cue south
South/north
Cue south
Cue south
South/north
Cue south
EG2_6-18_TT4_C7
24
FR (Hz)
0
EG2_7-26_TT4_C3
15
FR (Hz)
0
EG2_7-8_TT8_C4
14
FR (Hz)
0
EG3_8-15_TT1_C10
8
FR (Hz)
0
EG2_7-12_TT8_C6
11
FR (Hz)
0
EG3_8-19_TT1_C8
13
FR (Hz)
0
EG2_7-16_TT4_C12
9
FR (Hz)
0
SS2_2-14_TT11_C4
10
FR (Hz)
0
EG2_7-22_TT6_C3
11
FR (Hz)
0
SS2_3-8_TT12_C1
17
FR (Hz)
0

**Supplemental Figure 12. MEC/PaS non-grid spatial cell examples.** Path and spike plots (*top row*) and 2D allocentric firing rate maps (*bottom row*) for five MEC/PaS non-grid spatial cells recorded in the counterclockwise rotation session, five recorded in the clockwise rotation session, and ten recorded in the cue duplication session. Note that non-grid spatial cell representations were unchanged by the cue manipulations.

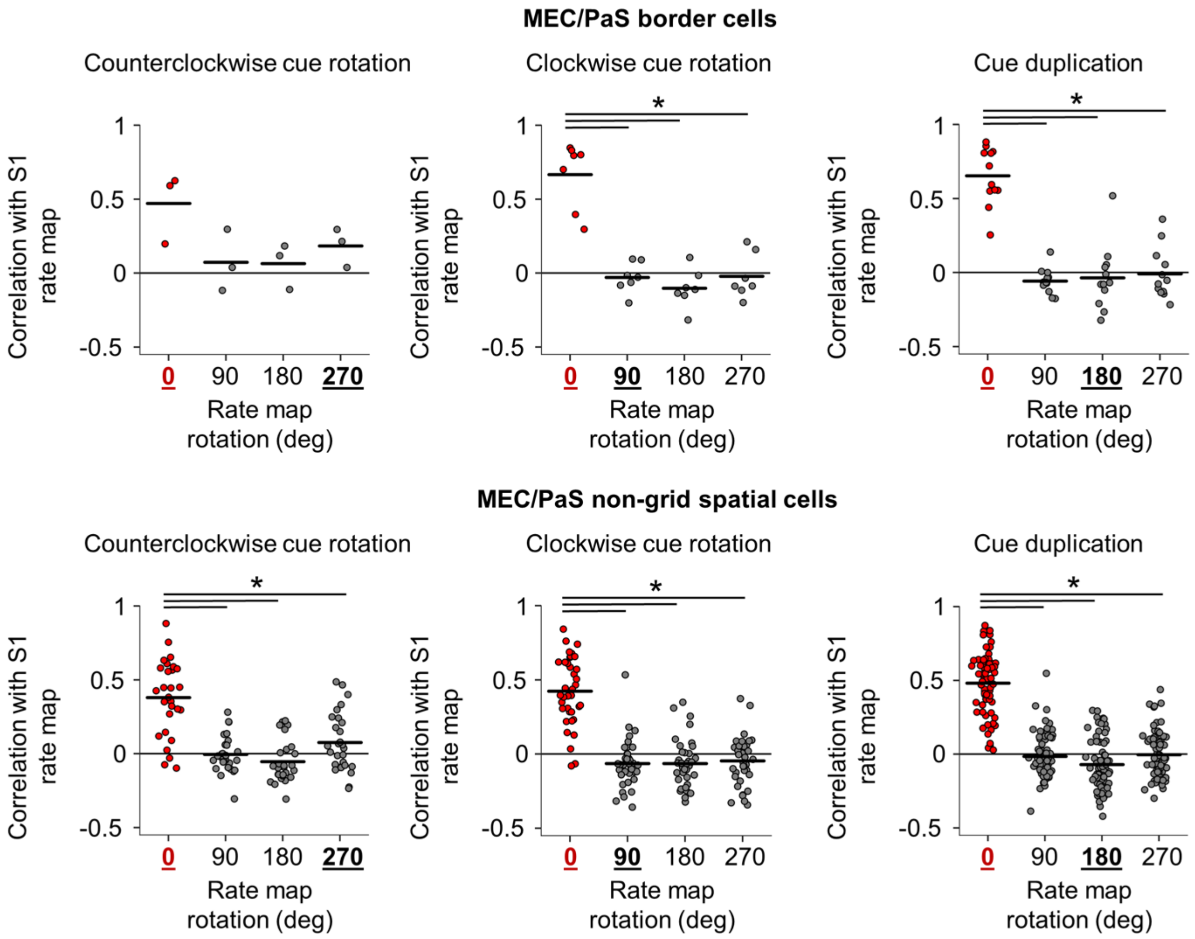


**Supplemental Figure 13. Rate map rotations for MEC/PaS border and non-grid spatial cells.** Strip plots showing correlations between the initial standard session rate map and the cue manipulation session rate map that had been rotated in 90° increments, for counterclockwise rotation sessions (*left*), clockwise rotation sessions (*middle*), and cue duplication sessions (*right*) for all MEC/PaS border cells (*top*) and non-grid spatial cells (*bottom*). Note that neither border cells nor non-grid spatial cells appeared to rotate their rate maps in the cue rotation session, though the small number of border cells in the counterclockwise rotation session (n = 3) did not allow for detection of statistical differences (border cells: counterclockwise rotation: repeated measures ANOVA, $F(3, 6) = 3.45$, $P = 0.092$; clockwise rotation: repeated measures ANOVA, $F(3, 18) = 29.05$, $P = 6.67e-4$; paired $t$-tests, 0° vs. 90°, $t(6) = 5.89$, $P = 0.0032$; 0° vs. 180°, $t(6) = 6.15$, $P = 0.0025$; 0° vs. 270°, $t(6) = 4.93$, $P = 0.0079$; cue duplication: repeated measures ANOVA, $F(3, 33) = 40.65$, $P = 2.05e-8$; paired $t$-tests, 0° vs. 90°, $t(11) = 12.79$, $P = 1.80e-7$; 0° vs. 180°, $t(11) = 7.16$, $P = 5.53e-5$; 0° vs. 270°, $t(11) = 7.89$, $P = 2.23e-5$; non-grid spatial cells: counterclockwise rotation: repeated measures ANOVA, $F(3, 81) = 29.00$, $P = 2.10e-9$; paired $t$-tests, 0° vs. 90°, $t(27) = 7.01$, $P = 4.67e-7$; 0° vs. 180°, $t(27) = 8.60$, $P = 9.65e-9$; 0° vs. 270°, $t(27) = 4.30$, $P = 5.98e-4$; clockwise rotation: repeated measures ANOVA, $F(3, 108) = 63.96$, $P = 6.16e-19$; paired $t$-tests, 0°

vs. 90°, $t(36) = 9.66$, $P = 4.63e\text{-}11$; 0° vs. 180°, $t(36) = 10.01$, $P = 1.80e\text{-}11$; 0° vs. 270°, $t(36) = 9.43$, $P = 8.72e\text{-}11$; cue duplication: repeated measures ANOVA, $F(3, 219) = 186.66$, $P = 2.68e\text{-}50$; paired *t*-tests, 0° vs. 90°, $t(73) = 16.56$, $P = 6.04e\text{-}26$; 0° vs. 180°, $t(73) = 17.57$, $P = 1.85e\text{-}27$; 0° vs. 270°, $t(73) = 16.94$, $P = 1.59e\text{-}26$).

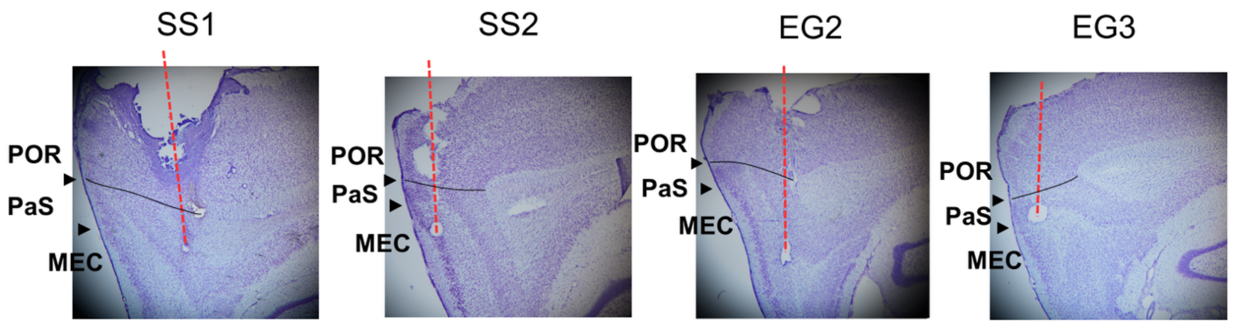


**Supplemental Figure 14. Histology.** Nissl-stained sagittal sections for all four rats showing electrode trajectories (red dotted lines) as well as estimated delineations between POR, PaS, and MEC (black triangles). An estimate of the location of the ventral POR border is indicated by a curved black line.
